# Multimodality Molecular Profiling Nominates Targetable Mechanisms in Progressive RV Dysfunction

**DOI:** 10.64898/2026.03.09.710504

**Authors:** Jenna B. Mendelson, Jacob Sternbach, Minwoo Kim, Rashmi M. Raveendran, Ryan A Moon, Lynn M. Hartweck, Walt Tollison, John P. Carney, Todd Markowski, LeeAnn Higgins, Sally. E. Prins, Felipe Kazmirczak, Kurt W. Prins

## Abstract

**Background:** Right ventricular dysfunction (RVD) is a robust predictor of mortality in multiple cardiovascular diseases. Currently, it remains unclear whether the severity of RVD corresponds to distinct cellular and molecular alterations, and this has important implications for defining optimal therapeutic targets. To address this knowledge gap, we performed a multi-omics evaluation of pulmonary artery banded (PAB) pigs with differing degrees of RV compromise.

**Methods:** PAB pigs were stratified into mild and severe RVD groups using an RV ejection fraction cutoff of 35%. RV tissue from control, mild RVD, and severe RVD animals was analyzed using single-nucleus RNA sequencing, mitochondrial and cytoplasmic proteomics, and phosphoproteomics. Histological analyses corroborated multi-omic findings.

**Results:** Cardiac MRI revealed progressive structural and functional alterations in mild and severe RVD pigs. snRNAseq demonstrated that advancing RVD was associated with loss of cardiomyocytes, accumulation of efferocytosis-impaired macrophages, and dysregulated endothelial cells and pericytes. Combined transcriptomic and proteomic analyses showed escalating impairments of complex cardiomyocyte metabolism with worsening RVD. RV microvasculature was compromised with severe RVD as there were alterations in endothelial cell/pericyte genetic regulation, co-localization patterns in RV sections, and ectopic cardiomyocyte HIF1 expression. Analysis of both mitochondrial and global proteostasis revealed greater compromise in mitochondrial proteostasis, including downregulation of mitochondrial proteases, chaperones, and ribosomes. Paradoxically, cytoplasmic ribosomes were upregulated in severe RVD. The predicted kinome and phosphatome were uniquely altered in mild RVD as compared to severe RVD. Finally, integration of multi-omic approaches identified insufficient mitochondrial unfolded protein response, impaired macrophage efferocytosis, and activation of the ribotoxic stress response as potential contributors to severe RVD.

**Conclusions:** Our multi-omic analysis defines the cellular and molecular landscape of progressive RVD and nominates druggable pathways that may promote progressive RV dysfunction. Future studies are needed to determine how targeting these pathways influences RV phenotypes.

## Introduction

Cardiovascular diseases (CVDs) remain the number one cause of death worldwide^1^. In numerous CVDs, ranging from systolic and diastolic heart failure^2^, valvular cardiomyopathy^3^, and congenital heart disease^4,5^, the presence of right ventricular dysfunction (RVD) predicts adverse outcomes. While the mechanisms underlying RV compromise are most extensively studied in pulmonary arterial hypertension^6^, these results may be pertinent for other diseased states, as RVD frequently occurs in the setting of heightened afterload^7^. At present, there are no effective approaches that directly augment RV function. Therefore, defining the pathological mechanisms in RV pressure overload may help nominate new therapies for several CVDs.

RVD is frequently dichotomized into either a compensated or decompensated state based on the severity of systolic defects, extent of RV dilation/hypertrophy, and how these physiological phenotypes predict outcomes^8–10^. Researchers have nominated multiple pathways ranging from inflammation, metabolic derangements, disrupted cellular signaling, and alterations in proteostasis as contributors to RVD in both the compensated and decompensated states using rodent, large animal^11–13^, and human data^8,9^. However, an integrated, multimodal assessment of molecular phenotypes across the spectrum of RVD is lacking. This strategy has value as it could provide both critical insight into disease progression and enable the identification of novel therapeutic targets.

Here, we completed a comprehensive, multi-omic analysis of pulmonary artery banded pigs with varying degrees of RVD to define the cellular landscape and molecular mediators of progressive RVD. We compared control pigs to pigs with mild RVD (RVEF 35-43%) and pigs with severe RVD (RVEF<30%) as determined by cardiac MRI. We used single-nucleus RNA sequencing (snRNAseq) to define cell-specific alterations across the spectrum of progressive RV dysfunction. Then, we integrated these data with three distinct RV proteomic analyses to systemically define the molecular contributors to RVD. Finally, we highlight underexplored, druggable pathways that could be targeted using existing and potentially repurposable drugs to counteract severe RVD.

## Materials and Methods

### Animal Studies

Eleven Yorkshire castrated male pigs aged 29-73 days old and weighing 7.5±2 kg on the day of surgery were used in this study. Five castrated pigs of a similar age and weight were housed at the University of Minnesota Research Animal Resources Facility for six weeks, and served as controls. Pulmonary artery banding was performed as previously described^11–14^. Mild and severe RVD groups were determined using RV ejection fraction (control, n=*5*: 58±3%, Mild RVD, *n*=5: 38±4%, Severe RVD, *n*=6: 20±8%). Six weeks after pulmonary artery banding, we performed right heart catheterization and a cardiac MRI study as noted before^12^. After the MRI scan, animals were humanely euthanized for tissue collection. All cardiac MRI images were blindly analyzed by FK. Experiments were approved by the University of Minnesota IACUC.

### RV histology analysis

Cardiomyocyte cross-sectional area was evaluated using paraffin embedded RV tissue sections stained with AlexaFluor-488 conjugated wheat germ agglutinin (WGA-488). Sections were stained with isolectin-B4 (IB4) to analyze capillary density. Fibrosis of RV free wall sections was performed to evaluate total section fibrosis and peri-vascular fibrosis. For peri-vascular fibrosis, images were collected and the amount of fibrosis per vessel was determined. Images were analyzed with FIJI by scientists blinded to treatment groups.

### Single nucleus RNAsequencing (snRNAseq) analysis

Nuclei were isolated from four pigs per experimental group at the same time following a protocol for flash frozen cardiac tissue^12,15,16^. Nuclei were then stained with propidium iodide and purified with fluorescence activated cell sorting (FACS) at the University of Minnesota Flow Cytometry Resource. The University of Minnesota Genomics Center completed library preparation, sequencing, and alignment to the pig genome (Sscrofa10.2). snRNAseq analysis was performed in RStudio v4.4 through the Minnesota Supercomputing Institute using Seurat v5^17^. Data filtering, normalization, scaling, dimensionality reduction, and clustering were completed using our established workflow^16,17^. We identified 14 clusters and assigned cell types at the cluster level using the FindConservedMarkers function in Seurat. We determined cell identities using the Single Cell Portal as previously described^16,18^. We found 3 clusters that did not contain genes consistent with any cell type, which were concluded to be noise and removed from further analysis. Correlation analysis between relative abundance and ejection fraction was completed with Metaboanalyst^19^. DESeq2^20^ was used for gene expression analysis of pseudobulked nuclei. Genes with |log_2_FoldChange| ≥ 0.5 and an adjusted *p*-value < 0.05 were considered differentially expressed. Due to the high number of differentially expressed cardiomyocyte genes when comparing severe to control and severe to mild, k-means clustering (k=3) was performed before pathway analysis using STRING v12.0^21^. However, more than three unconnected groups were identified, and most genes were in cluster 1, which was used for pathway analysis using the KEGG database in ShinyGO v0.82^22^. We removed the following cardiomyocyte genes from non-cardiomyocyte cell types: natriuretic peptide B (NPPB), troponin type 1 (TNNT1), desmin (DES), beta myosin heavy chain (MYH7), troponin C1 (TNNC1), myosin light chain (MYL2), and the top 100 genes expressed in cardiomyocytes^12,16,23^. Module scores were computed with the AddModuleScore function in Seurat to assess cell specific pathway activity after pseudobulking. Gene sets of pathways of interest were compiled using MitoCarta^24^ and KEGG. We employed the CellChat V2 package to evaluate predicted cell-cell communication^25^ from macrophages. Macrophage and fibroblast subclusters were determined after initial cell identification clustering using a resolution that identified subgroups with distinct gene expression. After subsetting macrophages, DEG analysis was performed using the FindMarkers function in Seurat with a Wilcoxon rank-sum test. Genes with |log2FoldChange| ≥ 0.5 and an adjusted p-value < 0.05 were considered differentially expressed and used for subsequent pathway analysis using the KEGG database on ShinyGO v0.82^22^.

### RV Mitochondrial and Cytosolic Proteomics

RV mitochondrial enrichments and the corresponding cytosolic fraction from all 16 pigs used in this study were isolated with a mitochondrial isolation kit (Abcam, Cambridge, MA) for TMT 16-plex proteomics analysis at the University of Minnesota Center for Metabolomics and Proteomics^13,26,27^. Global proteomics changes were assessed using hierarchical cluster analysis and ANOVA in Metaboanalyst^19^. Proteins with an ANOVA adjusted *p*-value cutoff < 0.1 were selected for pathway analysis. A pathway score was calculated by summing the abundance of each protein in the pathway and dividing this sum by the average of the control samples to assess relative protein abundance of the whole pathway. Protein sets were assembled from MitoCarta^24^ and KEGG databases.

### Cell-Specific Confocal Microscopy Analysis

Flash frozen RV samples embedded in optimal cutting temperature compound (OCT)^12^ from the pigs used in the snRNAseq analysis were used for immunohistochemistry analysis. To identify macrophages, OCT-embedded tissue slices were stained with primary antibody to galectin 3 (Abcam, AB76245) at a 1:50 dilution following our standard procedure^12,16^. Images were obtained on a Zeiss LSM900 Airyscan 2 confocal microscope, and the number of macrophages in each field was counted by a blinded reviewer. Pericyte-endothelial cell colocalization was calculated by staining OCT-embedded RV tissue sections with primary antibody to platelet-derived growth factor receptor beta (PDGFRβ, Cell Signaling Technology, 3169S) at a 1:50 dilution. Images were obtained on a Zeiss LSM900 Airyscan 2 confocal microscope, and pericyte-endothelial localization was blindly quantified using Zen Software.

### Phosphoproteomics and Phosphatome Analysis

Flash frozen RV samples were enriched for phosphorylated peptides and analyzed by mass spectrometry at the University of Minnesota Center for Metabolomics and Proteomics^28^. Sequenced proteins with no identified phosphorylation sites were removed. Proteins that were significantly (adjusted *p*-value<0.05, Proteome Discoverer Software) elevated or reduced in mild or severe RVD compared to control were selected for kinome enrichment analysis using Kinase Enrichment Analysis v3^29^. The top 10 kinases identified by MeanRank score using Kinase Enrichment Analysis were visualized on kinome maps using Coral^30^. Changes in the phosphatome were evaluated using both the cytoplasmic and mitochondrial fractions. Differentially expressed proteins were defined with a false-discovery rate of 0.1 when comparing the three experimental groups. Post-hoc analysis defined the direction of change, and phosphatases were visualized on a phosphatome map using CoralP^31^.

### Statistical Analysis

When multiple measurements were obtained from a single animal (i.e., multiple cells measured from the same RV), the mean per animal was used for subsequent statistical analysis. Data normality was evaluated using the Shapiro-Wilk test. To compare the means of three groups, one-way analysis of variance (ANOVA) with Tukey post-hoc analysis was used when the data were normally distributed. If the data were not normally distributed, the Kruskal-Wallis ANOVA with Dunn’s post-hoc analysis was used. When comparing the means of two groups, the unpaired t-test was used if the data were normally distributed. The Mann-Whitney U test was performed if the data were not normally distributed. Statistical significance was defined as *p*-value <0.05. Statistical analyses and graphing were performed with GraphPad Prism version 10. Data are presented as mean ± standard deviation if normally distributed or median and interquartile range if not normally distributed. Graphs show the mean or median, along with all individual values. All values were included in the statistical analysis. No experiment-wide multiple test corrections were applied.

### Selection of Representative Images

Immunostaining was determined to be specific if it was distinct from staining with secondary antibody alone, and non-specific antibodies. Representative images were selected based on quality, and those images that most closely reflected the mean/median data.

### Data availability

Data sets, analysis, and study materials will be made available upon request to other researchers for the purpose of the results or replicating the procedures. All data from omics experiments are publicly available. The snRNAseq raw FASTQ files are available at the NCBI Gene Expression Omnibus. The mass spectrometry proteomics and phosphoproteomics data are available at Figshare: doi: 10.6084/m9.figshare.25681464.

## Results

### Cardiac MRI and Histological Analysis Defined Progressive RV Remodeling and Dysfunction in Pulmonary Artery Banded Pigs

We performed comprehensive cardiac MRI, hemodynamic, and histological assessment of animals from the three experimental groups: control (*n*=5), mild RV dysfunction (Mild RVD, *n*=5), and severe RV dysfunction (Severe RVD, *n*=6). Cardiac MRI revealed a stepwise decline in RV ejection fraction (control: 58±3%, Mild RVD: 38±4%, Severe RVD: 20±8%, **Figure 1A-B**), RV-pulmonary artery (PA) coupling (**Figure 1C**), and an accompanied increase in RV dilation (**Figure 1D**), tricuspid regurgitation (**Figure 1E**), and RV mass (**Figure 1F**). Interestingly, RV cardiomyocyte hypertrophy was comparable between mild and severe RVD animals (**Figure 1G**). Hemodynamic characterization demonstrated elevated RV afterload as RV function deteriorated. Finally, magnetic resonance angiography revealed pulmonary artery stenosis was similar between groups subjected to PAB, but slightly higher in the severe RVD group (**Figure 1H-I**).

**Figure 1:**
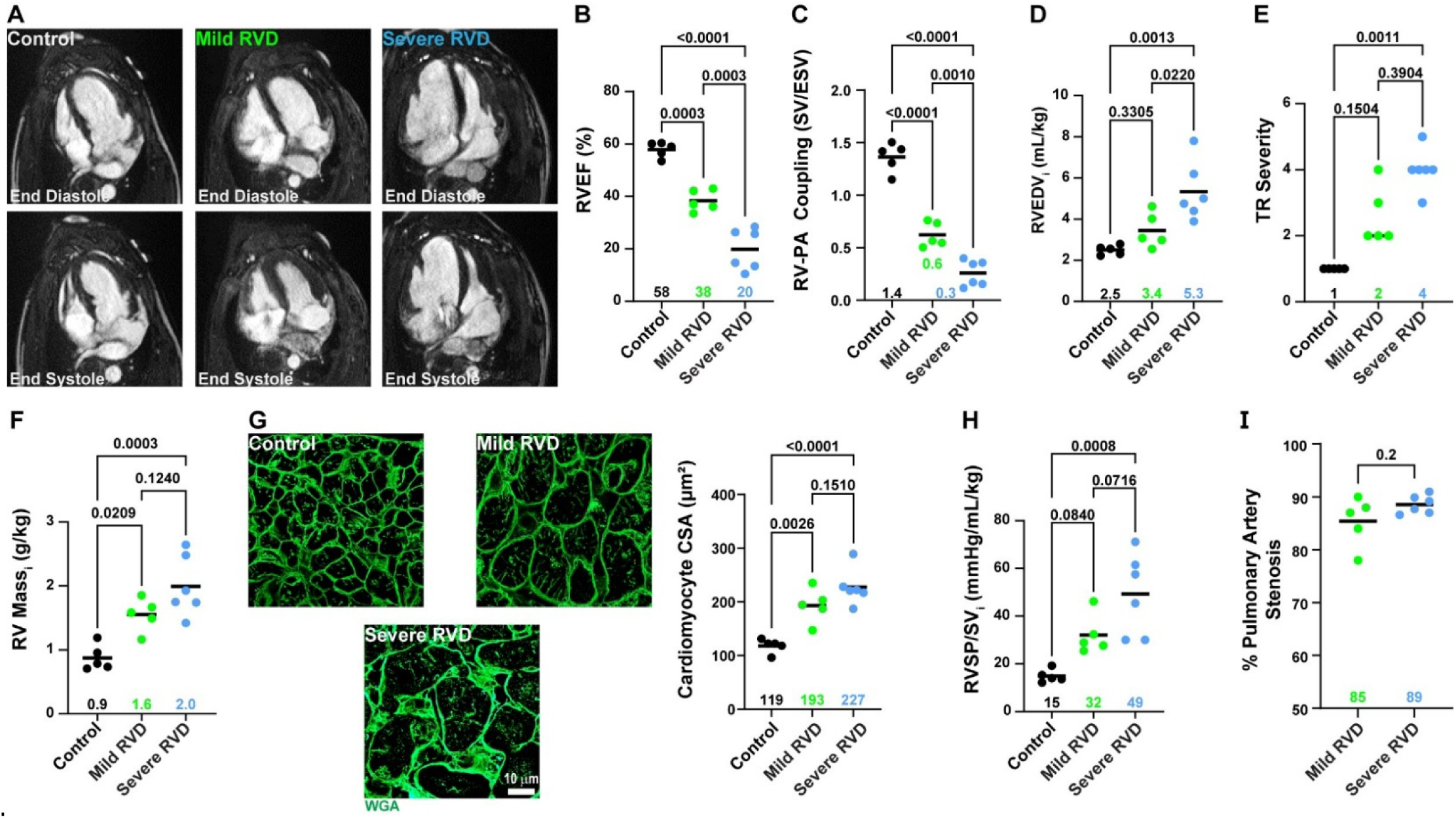
Physiological and histological characterization of control, mild RVD, and severe RVD pigs. (A) Representative four-chamber cMRI images. (B) RV function was significantly reduced in mild RVD and lowest in severe RVD (control: 58±3% *n*=5, mild RVD: 38±4% *n*=5, severe RVD: 20±8% *n*=6, one way ANOVA with Tukey’s post-hoc, means shown). (C) RV-PA coupling dropped as RV function worsened (control: 1.4±0.1, mild RVD: 0.6±0.1, severe RVD: 0.3±0.1, one way ANOVA with Tukey’s post-hoc, means shown). (D) RV dilation (control: 2.5±0.2 mL/kg, mild RVD: 3.4±0.8 mL/kg, severe RVD: 5.3±1.4 mL/kg, one way ANOVA with Tukey’s post-hoc, means shown), (E) tricuspid regurgitation (control: 1 [1,1], mild RVD: 2.0 [2,3.5], severe RVD: 4.0 [3.8,4.3], Kruskal-Wallis test used with Dunn’s post-hoc, medians shown), and (F) RV mass increased with declining RV function (control: 0.9±0.2 g/kg, mild RVD: 1.6±0.3 g/kg, severe RVD: 2.0±0.5 g/kg, one way ANOVA with Tukey’s post-hoc, means shown). (G) Representative confocal microscopy images of RV sections stained with wheat germ agglutinin (green: WGA) to evaluate cardiomyocyte hypertrophy, which was similar between mild and severe RVD (control: 118±13 µm^2^, mild RVD: 193±32 µm^2^, severe RVD: 227±33 µm^2^, one way ANOVA with Tukey’s post-hoc, means shown). (H) RV afterload was elevated in both mild and severe RVD, and highest in severe RVD (control: 15±3 mmHg/mL/kg, mild RVD: 32±8 mmHg/mL/kg, severe RVD: 49±17 mmHg/mL/kg, one way ANOVA with Tukey’s post-hoc, means shown). (I) Pulmonary artery stenosis was comparable between mild and severe RVD (mild RVD: 85±5%, severe RVD: 89±2%, unpaired t-test, means shown)

### Single nucleus RNA sequencing delineated the cellular landscape of RV dysfunction

Next, we used snRNAseq (*n*=4 RVs/group) to define cell-specific changes in both relative abundances and gene activation patterns across the three experimental groups. After quality control, 226,750 nuclei were analyzed and we identified seven cell types: cardiomyocytes, macrophages, fibroblasts, endothelial cells, pericytes, neuronal cells, and lymphocytes (**Figure 2A-B, Supplemental Table 1**). The relative abundance of each cell type was comparable between the control and mild RVD groups (**Figure 2C-D**). In contrast, severe RVD was marked by a reduction in cardiomyocytes (control: 74.4%, mild RVD: 71.3%, severe RVD: 48.8%) and an expansion of noncardiomyocyte populations including macrophages (control: 4.9%, mild RVD: 7.6%, severe RVD: 12.7%), lymphocytes (control: 4.9%, mild RVD: 7.0%, severe RVD: 11.6%), pericytes (control: 1.8%, mild RVD: 2.5%, severe RVD: 5.9%), and endothelial cells (control: 1.3%, mild RVD: 1.7%, severe RVD: 6.3%, **Figure 2E**).

**Figure 2:**
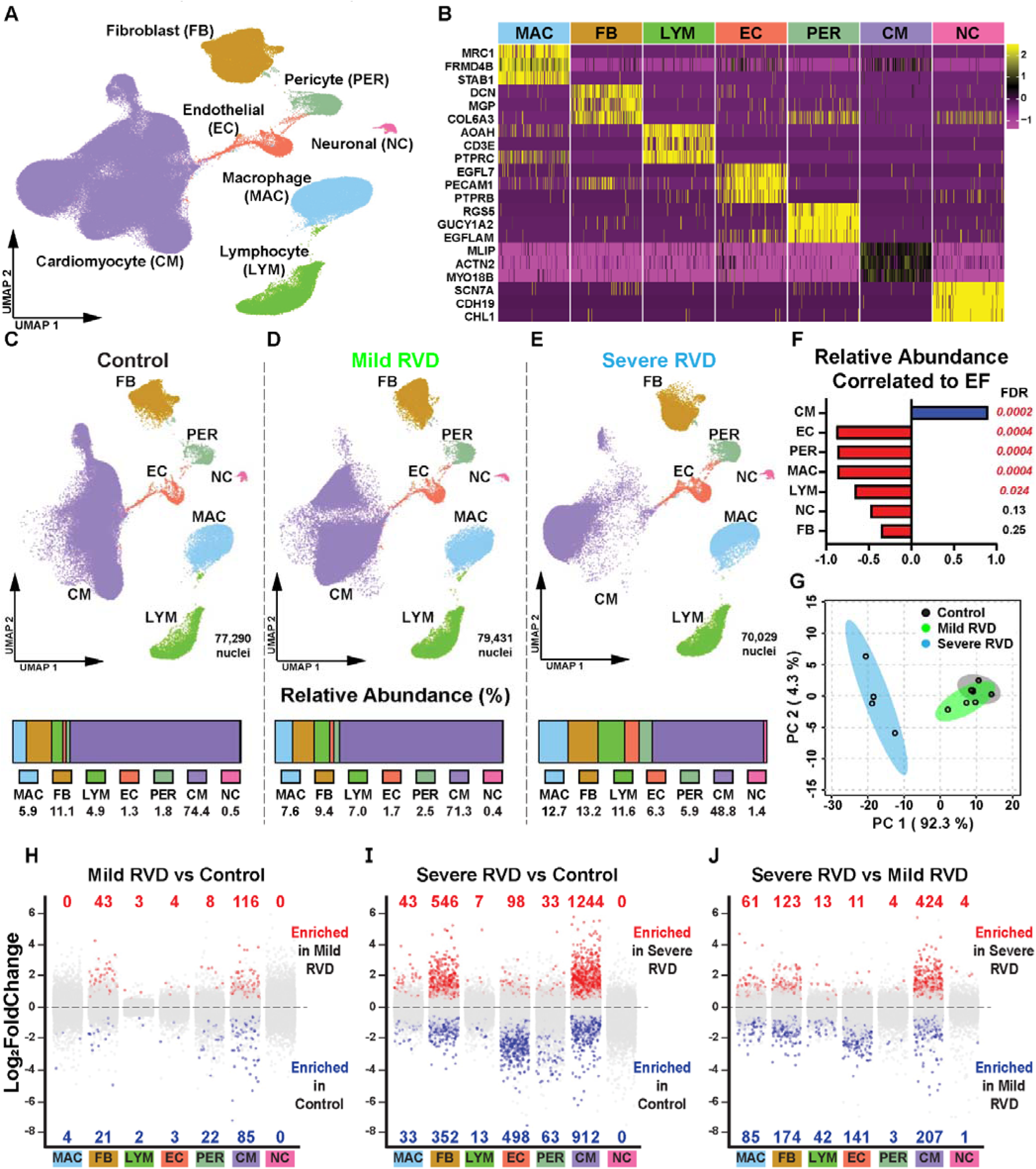
snRNAseq defined the changes in cellular composition with progressive RV compromise. (A) Uniform manifold approximation and projection (UMAP) visualization of the 7 cell types identified with snRNAseq. (B). Genes used to confirm cell identities. UMAP and cell type abundance of (C) control, (D) mild RVD, and (E) severe RVD. Cardiomyocyte abundance decreased with worsening RV function, whereas endothelial cell, pericyte, macrophage and lymphocyte abundances all increased. (F) Correlation analysis of relative abundance and ejection fraction (EF). (G) Principal component analysis distinguished experimental groups based on nuclei abundances. Pseudobulked differential expression analysis (red: Log2FoldChange > 0.5 and adjusted *p*-value <0.05, blue: Log2FoldChange < 0.5 and adjusted *p*-value < 0.05) comparing (H) mild RVD and control, (I) severe RVD and control, and (J) severe RVD and mild RVD. Cardiomyocytes were the cell type with the most differentially expressed genes regardless of the comparison, and overall, severe RVD vs control had the highest number of genes with altered expression.

Correlational analysis demonstrated only cardiomyocytes were positively correlated to RVEF, while endothelial cell, pericyte, macrophage, and lymphocyte abundances were negatively associated with RV function (**Figure 2F**). Principal component analysis of cellular composition revealed substantial overlap between control and mild RVD groups, but a marked shift in the severe RVD group (**Figure 2G**).

### snRNAseq characterized cell specific gene expression in RVD

We pseudobulked data from each cell type and identified the differentially expressed genes (DEGs) to define cell-type specific changes in gene expression (**Figure 2H-J**). We identified the most DEGs when analyzing severe RVD vs control. Across all cellular comparisons, cardiomyocytes had the most DEGs, followed by fibroblasts and endothelial cells. These data demonstrated substantial changes in the genetic landscape in multiple cells types with progressive RVD, and cardiomyocytes showed the most pronounced alterations.

### Cardiomyocyte Metabolic Derangements Corresponded with RV Impairment

To evaluate changes in cardiomyocyte gene regulation in an unbiased manner, we performed pathway analysis of the DEGs. In severe RVD, transcripts across multiple metabolic pathways, including oxidative phosphorylation, were broadly suppressed (**Supplemental Figure 1**). To complement these transcriptomic findings, we examined protein-level regulation using quantitative proteomics with mitochondrial and cytoplasmic enrichment strategies (**Supplemental Figure 2A**). Kyoto Encyclopedia of Genes and Genomes (KEGG) and Wiki pathway analyses of proteins that distinguished experimental groups highlighted widespread disruption of mitochondrial metabolic pathways in mitochondrial proteomics (**Supplemental Figure 2B-C**). However, altered pathways in the cytoplasm were not related to metabolism (**Supplemental Figure 2D-E**).

Because multiple metabolic pathways emerged from both unbiased transcriptomic and proteomic analyses, we systematically examined the regulation of oxidative phosphorylation, tricarboxylic acid (TCA) cycle, fatty acid oxidation, glycolysis, and ketone metabolism (**Figure 3A**). In cardiomyocytes, oxidative phosphorylation transcripts were suppressed in both mild RVD and severe RVD, whereas protein abundance was only diminished in severe RVD (**Figure 3B**). We identified a similar trend in the TCA cycle, although the reduction in the protein was not statistically significant (**Figure 3C**). Fatty acid oxidation transcript and protein levels were similar between the control and mild RVD groups, and lower in severe. In severe RVD, there was a nonsignificant decrease in cardiomyocyte transcript expression and a significant depletion in protein levels (**Figure 3D**). In the glycolysis pathway, transcript levels were lower only in severe RVD, but protein abundances were nonsignificantly increased in mild RVD (**Figure 3E**). In cardiomyocytes, transcript expression of the ketone metabolism pathway was similar between groups, and protein abundance was nonsignificantly reduced in severe RVD (**Figure 3F**). In summary, these data showed a progressive dysregulation of the transcripts and proteins involved in oxidative phosphorylation, TCA cycle, and fatty acid oxidation as RV function worsened (**Figure 3G**).

**Figure 3:**
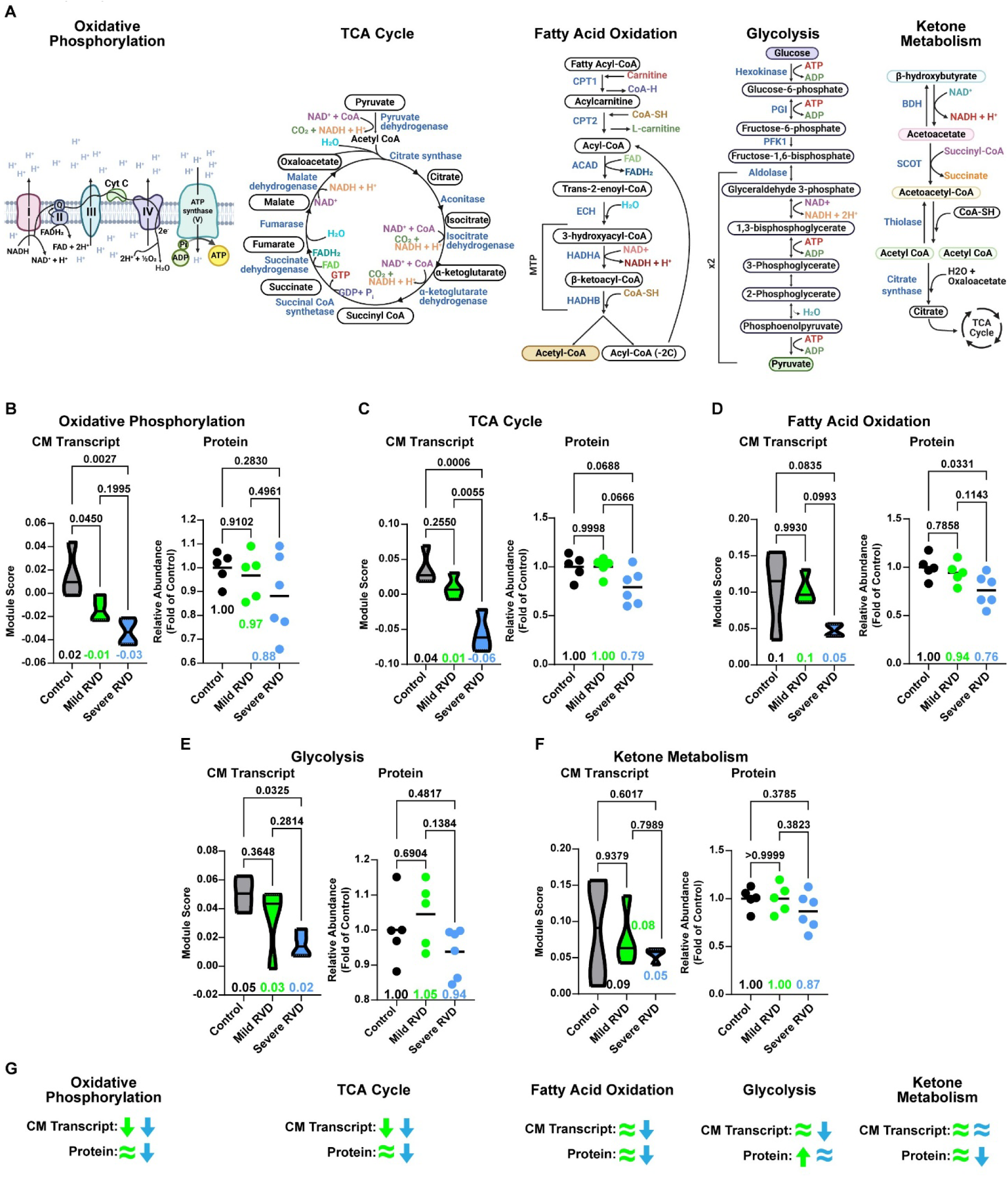
Disruptions in cardiomyocyte metabolism were observed in transcriptomic and proteomic data as RV function deteriorated. (A) Metabolic pathways selected for further evaluation. (B) Oxidative phosphorylation, (C) tricarboxylic acid (TCA) cycle, and (D) fatty acid oxidation module (transcriptomic) and pathway (proteomic) scores were reduced in severe RVD, but mild RVD protein abundances were similar to control. (E) Glycolysis transcripts were reduced in severe RVD, but protein abundance was similar to that of the control. (F) Ketone metabolism transcripts weren’t altered in severe RVD cardiomyocytes, but protein levels were reduced. (G) Summary of cardiomyocyte transcript expression and protein abundances of each pathway. Green (mild RVD) and blue (severe RVD) arrows and equal signs show changes in selected pathways compared to control. Transcriptomic data depicted as module score, proteomic data shown as total pathway abundance normalized to control. One way ANOVA with Tukey’s post-hoc, means shown in B-F. CM: cardiomyocyte.

### Disrupted Macrophage Efferocytosis Was Observed in Severe RVD

Because macrophage relative abundance was strongly associated with RVEF, we further characterized macrophages using snRNAseq data. Macrophages were first classified as either resident (LYVE1, MRC1, and CCL24 enriched) or recruited (CXCL10, PLAC8, and IFIT1 enriched^32^, **Figure 4-B**) based on gene expression profiles. Both mild and severe RVD pigs had a higher proportion of recruited macrophages (**Figure 4C**). Relative to all nuclei, severe RVD had more resident and recruited macrophages (**Figure 4D**). In agreement with our snRNAseq analysis, immunohistochemical analysis confirmed a progressive increase in macrophages in RV sections (**Figure 4E and F**).

**Figure 4:**
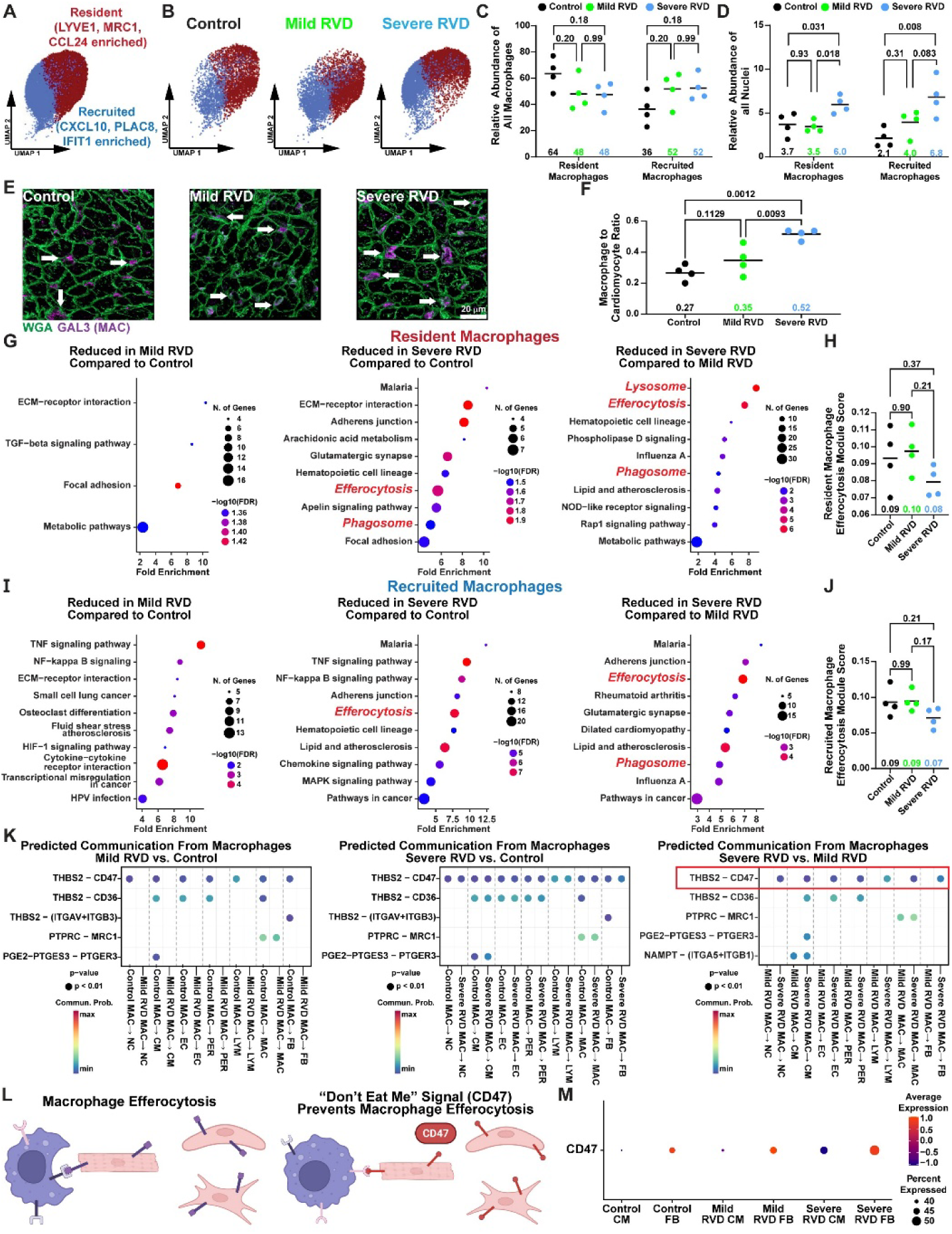
Severe RV dysfunction was associated with impaired macrophage efferocytosis. (A) UMAP visualization of macrophage subtypes based on resident or recruited marker gene expression. (B) UMAP of resident and recruited macrophages split by experimental group. (C) Mild and severe RVD pigs had a higher proportion of recruited macrophages (D) Severe RVD had more resident and recruited macrophages relative to all nuclei (E) Representative images of RV sections stained with galectin-3 (GAL3) demonstrated macrophage abundance was highest in severe RVD (white arrows: macrophages, green: WGA; purple: GAL3) as quantified (F) by the number of macrophages in each field averaged by the number of cardiomyocytes. (G) Pathway analysis of differentially expressed resident macrophage genes identified downregulation of efferocytosis, phagosome, and lysosome pathways in severe RVD. (H) Module score of resident macrophage efferocytosis was nonsignificantly reduced in severe RVD. (I) Efferocytosis and phagosome pathways are reduced in severe RVD recruited macrophages. (J) Efferocytosis module score was nonsignificantly reduced in recruited macrophages from severe RVD. (K) CellChat predicted the “don’t eat me” ligand cluster of differentiation 47 (CD47) only in severe RVD compared to mild RVD. (L) The presence of CD47 prevents macrophage efferocytosis. (M) CD47 expression was higher in fibroblasts and cardiomyocytes from the severe RVD group. LYVE1: lymphatic vessel endothelial hyaluronan receptor 1; MRC1: mannose receptor C type 1; CCL24: C-C motif chemokine ligand 24; CXCL10: C-X-C motif chemokine ligand 10, PLAC8: placenta associated 8; IFIT1: interferon-induced protein with tetratricopeptide repeats 1; CM: cardiomyocyte; FB: Fibroblast. One way ANOVA with Tukey’s post-hoc, means shown in C-D, F, H, J.

Pathway analysis determined how the genetic profiles of resident and recruited macrophages were altered. Metabolic pathways were enriched in severe RVD resident and recruited macrophages (**Supplemental Figure 3**). In particular, fructose and mannose metabolism were increased in resident macrophages, and cysteine and methionine metabolism were elevated in recruited macrophages. The pro-inflammatory pathways atherosclerosis, senescence, and AGE-RAGE signaling were also elevated in these macrophages.

The most striking alteration was impairments in macrophage efferocytosis. In resident macrophage, severe RVD animals exhibited downregulation of efferocytosis, phagosome, and lysosomal pathways as compared with both control and mild RVD animals (**Figure 4G**). Severe RVD animals showed a nonsignificant reduction in the efferocytosis module score, whereas the mild RVD animals had the same score as the controls (**Figure 4H**). In recruited macrophages, efferocytosis was impaired in severe RVD animals, as efferocytosis and phagosome pathways were suppressed when compared to control and mild RVD. In addition, there was a nonsignificant decrease in the efferocytosis module score in severe RVD (**Figure 4I-J**).

Then we used CellChat^25^ to predict differences in cellular communication from macrophages to other cell types across experimental groups, which again indicated altered macrophage efferocytosis in severe RVD. When communication predictions between severe RVD and mild RVD was compared, the thrombospondin-2 to cluster of differentiation 47 (CD47) communication was predicted only in severe RVD (**Figure 4K**). CD47 prevents macrophages from engulfing healthy cells^33^ (**Figure 4L**), but heightened CD47 expression can prevent efferocytosis of dead/dying pro-inflammatory cells. Fibroblasts and cardiomyocytes from the severe RVD group had higher CD47 expression than mild RVD or control (**Figure 4M**), which supported our CellChat findings. In total, these data suggested RV inflammation was heightened through dysregulated macrophage genetic programming via both ineffective efferocytosis and the induction of multiple pro-inflammatory pathways.

### Disruptions in RV Endothelial Cells and Pericytes Were Associated with RV Function

Next, we evaluated the cells of the RV vasculature because our snRNAseq data identified the abundances of endothelial cells and pericytes were significantly elevated in severe RVD (**Figure 5A-C**). Immunofluorescence validated a progressive increase in endothelial cell density in RV sections with worsening RV function (**Figure 5D-E**). In addition, there was reduced colocalization between capillaries and pericytes in severe RV sections (**Figure 5F-G**), suggesting altered pericyte localization. Finally, to assess a potential consequence of impaired RV microvascular function, we examined cardiomyocyte expression of hypoxia-inducible factor 1α (HIF1A). HIF1A transcripts were highest in severe RVD, consistent with impaired myocardial oxygen delivery (**Figure 5H**). Collectively, these findings indicate that despite increased vascular cell abundance in severe RVD, RV microvascular function may be compromised, leading to activation of hypoxia-responsive signaling in RV cardiomyocytes.

**Figure 5:**
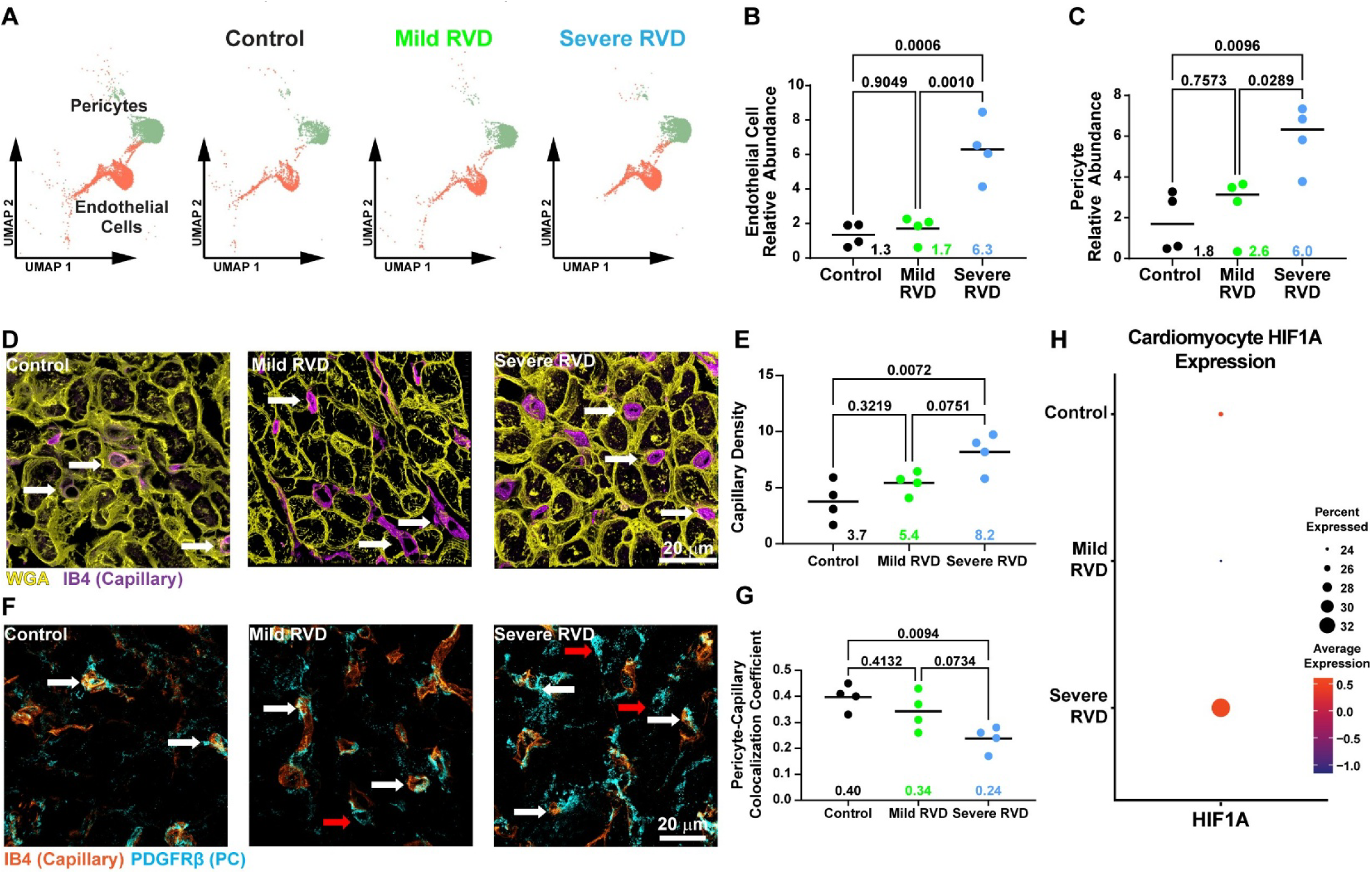
RV endothelial and pericyte abundances and colocalization patterns were associated with cardiomyocyte hypoxia signaling and with progressive RV compromise. (A) UMAP visualizing endothelial cells and pericytes in snRNAseq data. (B) Endothelial cell and (C) pericyte relative abundances were similar between mild RVD and control, but elevated in severe RVD. (D) Representative image of RV sections stained with isolectin B4 (IB4) showing capillary density is higher in severe RVD (white arrows: capillaries, yellow: WGA, purple: IB4), quantified as (E) the percent area occupied by endothelial cells. (F) Representative image of RV sections stained with IB4 and platelet derived growth factor receptor beta (PDGFRβ) to mark endothelial cells and pericytes, respectively. Endothelial cell-pericyte localization was altered in severe RVD (white arrows: co-localized endothelial cells and pericytes, red arrow: pericytes not located near endothelial cells, orange: IB4, blue: PDGFRβ) quantified with a colocalization coefficient. (H) Cardiomyocyte HIF1A expression was highest in severe RVD. One way ANOVA with Tukey’s post-hoc, means shown in B-C, E, G.

### Fibroblast Subcluster Analysis Identified a More Activated Fibroblast Phenotype in Severe RVD

Although the relative abundance of fibroblasts was similar between all 3 groups, their transcriptional profiles differed markedly. We identified three fibroblast subpopulations (**Supplemental Figure 4A**) with distinct distributions across experimental conditions (**Supplemental Figure 4B-C**). Nearly all fibroblast nuclei from control and mild RVD were in cluster 1, but fibroblasts from severe RVD predominantly occupied cluster 2. Cluster 2 fibroblasts represented an activated fibroblast state, based on elevated expression of fibronectin 1 (FN1), periostin (POSTN), and fibroblast activating protein^34^ (FAP, **Supplemental Figure 4D**). Despite these phenotypic changes, we did not observe significant changes in total RV fibrosis or perivascular fibrosis in RV sections (**Supplemental Figure 4E**).

### Distinct Proteostasis Phenotypes Were Associated with Severe RVD

Integrated proteomic analyses identified multiple proteostasis components as distinguishing features among the three experimental groups (**Supplemental Figure 2E**). Therefore, we systematically evaluated cytoplasmic and mitochondrial proteostasis pathways using combined proteomic and transcriptomic approaches. We first focused on mitochondrial proteostasis, examining pathways related to mitochondrial ribosomes (**Figure 6A**), import proteins (**Figure 6B)**, chaperones (**Figure 6C**), proteases (**Figure 6D**), the mitochondrial unfolded protein response (**Figure 6E)**, and mitophagy (**Figure 6F**). Cardiomyocyte transcript levels of mitochondrial ribosome, protein import, and chaperone pathways were significantly reduced in severe RVD compared to control. Module scores of mitochondrial proteases and the mitochondrial unfolded protein response (mitoUPR) were nonsignificantly reduced, while mitophagy pathway transcripts were slightly elevated in severe RVD. At the protein level, all mitochondrial proteostasis pathways showed reduced abundance in severe RVD, although these differences did not always reach statistical significance. In contrast, protein abundances in mild RVD were comparable to controls (**Figure 6G**), indicating that disruption of mitochondrial proteostasis was a feature specific to severe RVD.

**Figure 6:**
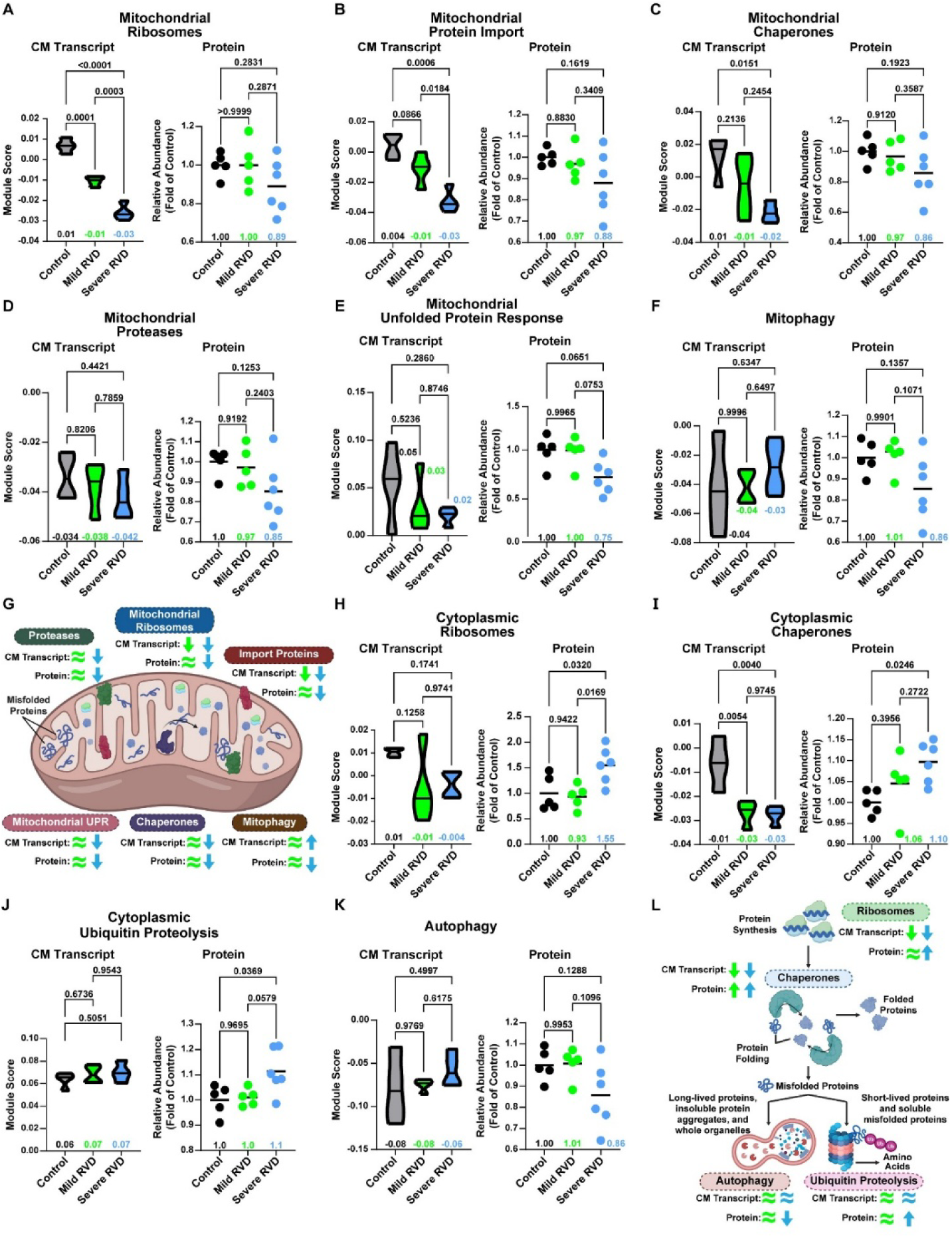
Global and mitochondrial proteostasis pathways were dysregulated in RVD, but mitochondrial proteostasis was most robustly impaired. (A) Cardiomyocyte mitochondrial ribosome and (B) protein import transcripts were reduced in mild RVD and severe RVD, but the proteins were only lowered in severe RVD. Both transcripts and proteins from (C) mitochondrial chaperones, (D) mitochondrial proteases and the (E) unfolded protein response were suppressed in severe RVD. (F) Cardiomyocyte mitophagy transcripts were increased in severe RVD, but reduced at the protein level. (G) Summary of cardiomyocyte transcript expression and protein abundances in each mitochondrial proteostasis pathway. Green (mild RVD) and blue (severe RVD) arrows and equal signs show changes compared to control. Cardiomyocyte gene expression of (H) Cytoplasmic ribosome and (I) chaperone pathways was decreased in mild and severe RVD, but protein abundance was elevated in severe RVD. (J) Ubiquitin proteolysis transcript expression was similar between groups, and protein was increased in severe RVD. (K) Autophagy pathway cardiomyocyte gene expression was not changed in mild or severe RVD, but protein abundance was reduced in severe RVD. (L) Summary of cardiomyocyte transcript expression and protein abundances of cytoplasmic proteostasis pathways. Green (mild RVD) and blue (severe RVD) arrows and equal signs show changes compared to control. Transcriptomic data depicted as module score; proteomic data shown as total pathway abundance normalized to control. One way ANOVA with Tukey’s post-hoc, means shown in A-F, H-K. CM: cardiomyocyte.

Then we probed our snRNAseq data and cytoplasmic proteomics to assess cellular proteostasis by examining ribosomes (**Figure 6H**), chaperones (**Figure 6I**), the ubiquitin-proteasome pathway (**Figure 6J**), and autophagy-mediated protein degradation (**Figure 6K**). In cardiomyocytes, transcript levels of cytoplasmic ribosomes were modestly, but not significantly, reduced in both mild and severe RVD, whereas chaperone transcripts were significantly decreased in both groups. Ubiquitin-proteasome and autophagy transcripts were unchanged across groups. In contrast, our proteomics data showed divergent regulation. The protein abundances of cytoplasmic ribosome, chaperone, and ubiquitin proteolysis pathways were all elevated in severe RVD. Similar to the mitophagy pathway, autophagy proteins were reduced in severe RVD. Cytoplasmic ribosome, ubiquitin proteolysis, and autophagy proteins were the same in mild RVD as in the control, while chaperone protein abundance was slightly increased (**Figure 6L**). We also evaluated protein ubiquitination and found ubiquitinated protein abundance the abundance of ubiquitinated proteins was higher in mild and severe RVD than in controls (**Supplemental Figure 5)**. In summary, these data suggested global proteostasis was not as compromised as mitochondrial proteostasis, but highlighted impairments in autophagy and ubiquitin-mediated protein quality control.

### The Kinome and Phosphatome Were Restructured in PAB Pigs

Then, we attempted to delineate what signaling cascades contributed to compromised RV function by evaluating the kinome and phosphatome using our combined proteomics approaches. We first performed phosphoproteomics analysis of RV tissue from the three experimental groups (**Supplemental Figure 6A**). Hierarchical cluster analysis demonstrated alterations in phosphoprotein abundances in control, mild RVD, and severe RVD specimens (**Supplemental Figure 6B**). We then mapped kinases predicted to be dysregulated as RVD progressed. In mild RVD, the top ten kinases with increased predicted activity compared to controls were LRRK2, PDK3, AKT1, TBK1, RIOK1, RAF1, ILK, EGFR, PHKG2, and CDK4 (**Figure 7A**). Severe RVD showed a distinct kinase activation profile, with partial overlap with mild RVD. Kinases predicted to be activated in severe RVD included TTN, FYN, GSK3B, ABL1, LATS2, EGFR, MARK1, NTRK1, STK11, and AKT1 (**Figure 7B**).

**Figure 7:**
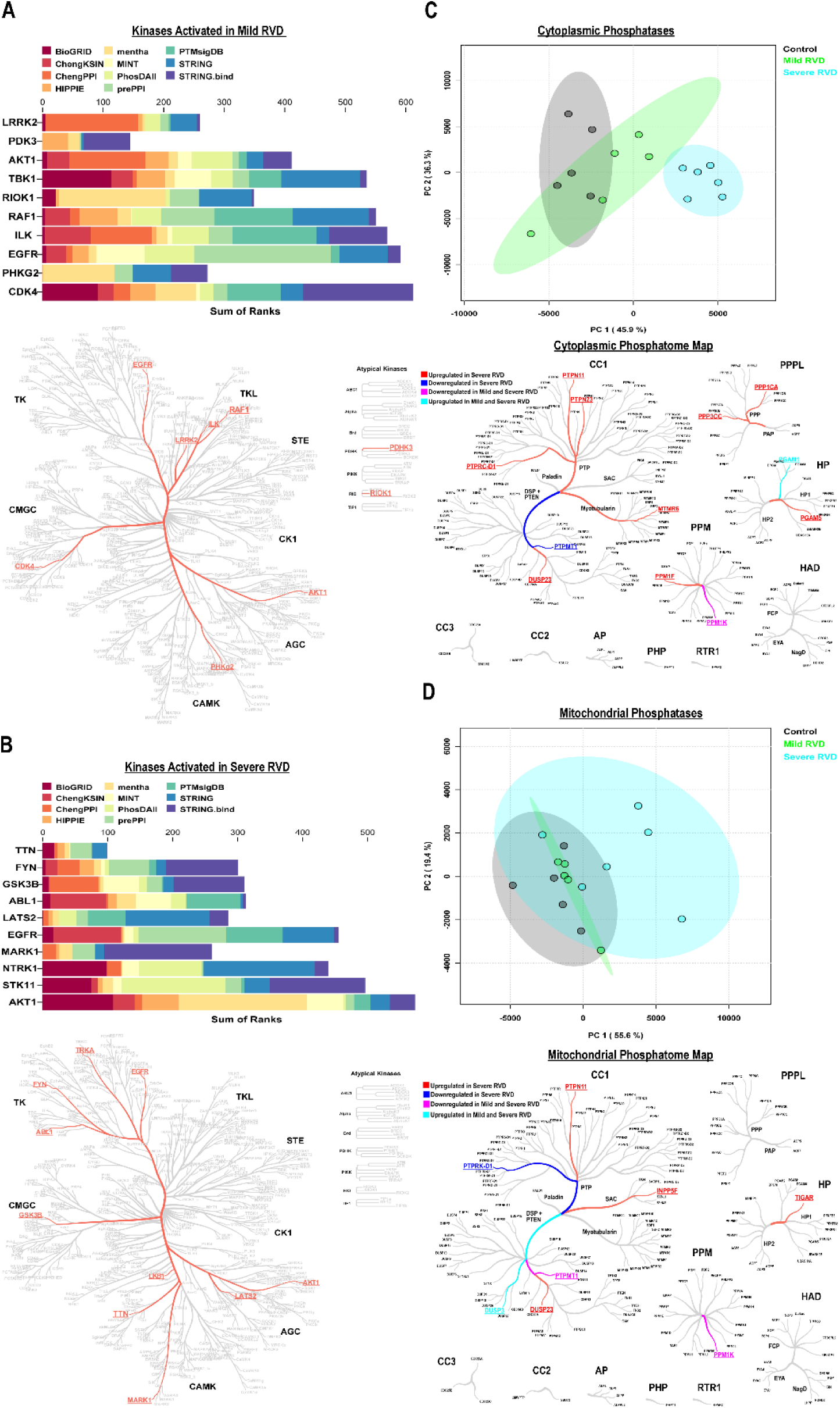
Distinct alterations in cellular signaling as predicted by kinome and phosphatome evaluations in RVD. Top 10 predicted kinases activated in (A) mild RVD and (B) severe RVD. (C) Principal component analysis and the phosphatase map showed severe RVD cytoplasmic phosphatases differ from those in control and mild RVD. (D) Principal component analysis and phosphatase map revealed significant overlap between control and mild RVD mitochondrial phosphatases, but severe RVD was more discrete.

To complement our kinome analysis, we examined phosphatome alterations in both the cytoplasmic and mitochondrial fractions. In the cytoplasmic fractions, principal component analysis demonstrated the severe RVD was distinct from control and mild RVD (**Figure 7C**). When we mapped the altered phosphatases, most were upregulated only in severe RVD. Nine phosphatases were uniquely upregulated in severe RVD, one was downregulated in severe RVD, one was upregulated in both mild and severe RVD, and one was downregulated in both groups (**Figure 7C**). In mitochondrial fractions, phosphatome changes were generally less pronounced (**Figure 7D**). Principal component analysis again revealed overlap between control and mild RVD, but severe RVD was more distinct. However, four phosphates were upregulated in the severe RVD, one was downregulated in the severe RVD, two were downregulated in both mild and severe RVD, and one was upregulated in both groups. Together, these findings suggested that phosphatome alterations, particularly in severe RVD, may contribute to the observed changes in protein phosphorylation and thus could play a role in altered intracellular signaling pathways.

### Integration of Multi-Omic Data Delineates New Druggable Targets

Finally, we integrated our multi-omic datasets to identify potential therapeutic targets for progressive RV failure (**Figure 8**). We first focused on the mitochondrial unfolded protein response (mitoUPR) because our snRNAseq data demonstrated mitoUPR transcripts were downregulated in the severe RVD group. In particular, the abundance of Lon peptidase 1 (LONP1), a critical component of the mitoUPR^35^, was only reduced in severe RVD. Meclizine, an available antihistamine used to treat motion sickness and vertigo, activates the mitoUPR^36^, suggesting it could potentially be repurposed to enhance mitochondrial protein quality control and RV metabolism.

**Figure 8:**
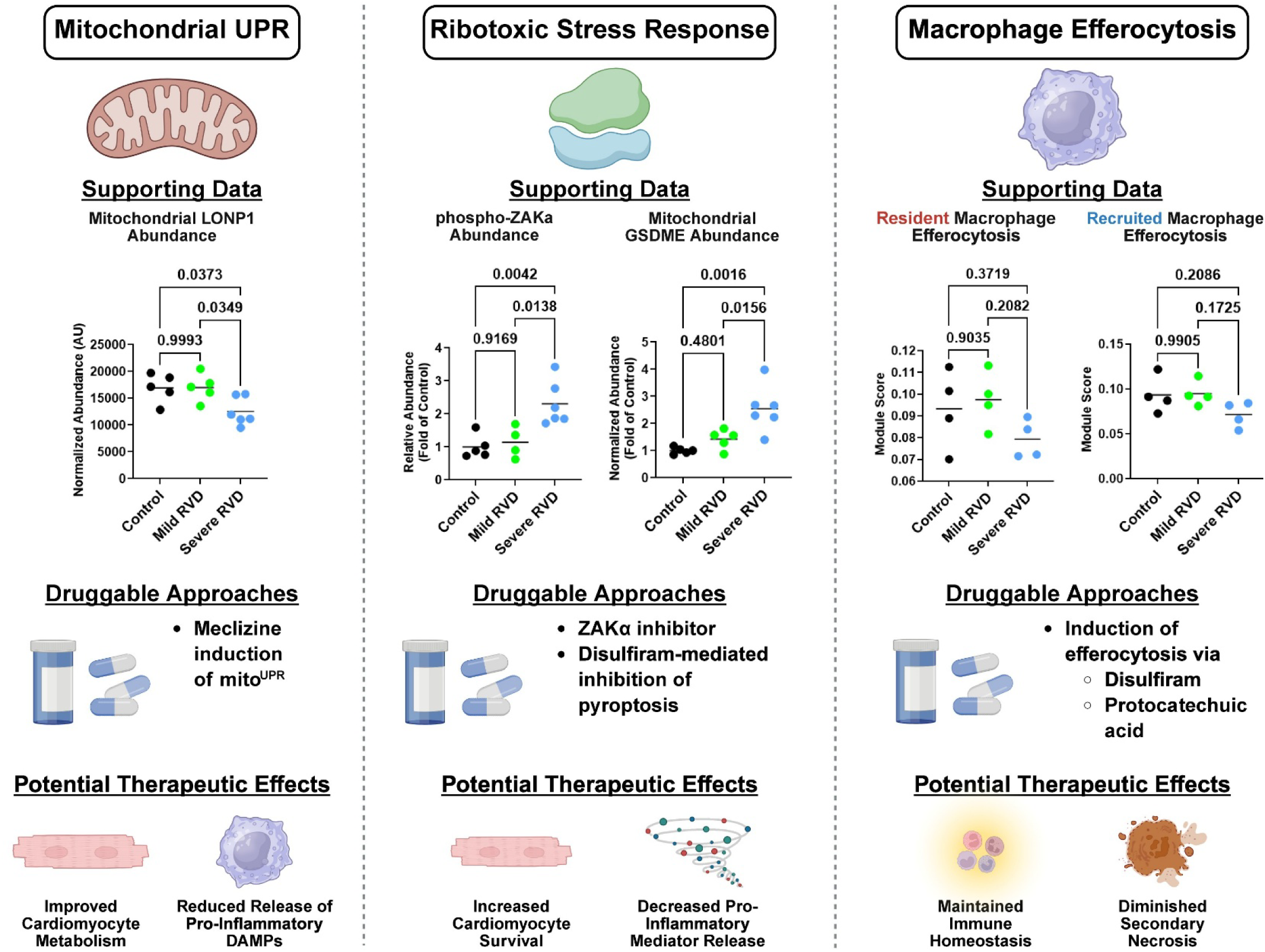
New mechanisms and proposed approaches to combat severe RV dysfunction. We proposed 3 new pathways with existing FDA approved therapies to target for RVD: (1) the mitochondrial unfolded protein response (mitoUPR), (2) the ribotoxic stress response, and (3) macrophage efferocytosis.

Next, we probed how the accumulation of ribosomes in severe RVD could be pathogenic by profiling components of the ribotoxic stress response. The ribotoxic stress response induces pyroptosis, an inflammatory type of cell death, via the signaling kinase ZAKα and promotion of gasdermin pore formation (**Supplemental Figure 7A**). Phosphoproteomics data revealed phospho-ZAKα levels were increased only in the severe RVD group (**Supplemental Figure 7B**). Additionally, gasdermin D and E abundances were higher in the mitochondrial fraction (**Supplemental Figure 7C**), which suggested gasdermin pores may be present in the severe RVD animals. The ribotoxic stress response can be directly inhibited by ZAK-N-1, a small-molecule ZAKα inhibitor^37^. Additionally, pyroptosis can be suppressed with disulfiram, an FDA-approved drug for alcoholism. Together, these findings highlight two potential strategies to counteract ribotoxic stress and the associated pyroptosis, to support RV function.

We also identified a deficit in macrophage efferocytosis that was specific to severe RVD as efferocytosis and lysosomal transcripts were reduced in both recruited and resident macrophages. Enhancing efferocytosis may facilitate the clearance of dying, pro-inflammatory cells from the RV to limit RV damage. Two currently approved medications, disulfiram and protocatechuic acid, are known to enhance macrophage efferocytosis^38,39^. Thus, these drugs could potentially be repurposed to improve macrophage clearance of pro-inflammatory cells to counteract RVD.

## Discussion

Here, we provide a comprehensive, multi-omic analysis of progressive RV dysfunction in male pulmonary artery banded pigs to delineate the cellular and molecular alterations along the spectrum of RV impairment. Data from snRNAseq and multiple proteomics approaches demonstrate reductions in several metabolic pathways, including fatty acid metabolism, oxidative phosphorylation, and the TCA cycle with worsening RV function. We propose impaired mitoUPR could contribute to the metabolic defects we observe in RVD as downregulation of mitoUPR pathways correlates with the reduction in metabolic enzymes. Our snRNAseq data identify an increase in the abundance of both resident and recruited macrophages as RV function worsens. A defining characteristic of these macrophages is impaired efferocytotic function. Integrated proteomics analysis reveals unique deficits in mitochondrial proteostasis but relatively preserved cellular proteostasis. There is an upregulation of cytoplasmic ribosomes, which may be necessary to support cardiomyocyte hypertrophy. However, higher levels of phosphorylated ZAKα suggest these ribosomes are functionally impaired and promote activation of the ribotoxic stress response. In summary, these data identify three understudied, but druggable, pathways that may promote severe RVD: (1) the mitochondrial unfolded protein response, (2) macrophage efferocytosis, and (3) ribosomal stress response.

We nominate the mitoUPR as a therapeutic target for RV failure and suggest it may contribute to the metabolic defects commonly described in RVD^8,9,12^. Protein accumulation in the mitochondria triggers the mitoUPR, which increases nuclear transcription of mitochondrial chaperones and proteases to refold or break down misfolded, dysfunctional proteins^40^. Under normal physiological conditions, mitochondrial protein quality control is crucial for mitochondrial metabolic function^41^. In addition, mitoUPR transcription factors induce the expression of several enzymes to enhance mitochondrial metabolic function even if mitochondria protein regulating capacity is reduced^41^. Moreover, the mitoUPR stimulates antioxidant gene transcription to mitigate reactive oxygen species (ROS) formation^42^, which is deleterious because ROS damages proteins^43^, activates cellular stress responses like the ribotoxic stress response^44^, and can initiate inflammatory forms of cell death such as ferroptosis and pyroptosis^45^. Thus, it is conceivable that impairments in the mitoUPR could contribute to metabolic-inflammatory dysregulation in RV failure. Although not evaluated in RVD, in preclinical models of LV pressure overload, the mitoUPR is downregulated and its stimulation reduces cardiomyocyte death and LV dysfunction^43^. Importantly, in humans with aortic stenosis induced pressure overload, mitoUPR activity correlates with tissue damage^43^, suggesting this pathway has direct relevance to human cardiac biology.

Another robust cellular signature of progressive RVD is the accumulation of efferocytosis-limited macrophages. The removal of dead and dying cells through efferocytosis is a primary function of macrophages to maintain tissue homeostasis^46^. Impaired efferocytosis contributes to cardiac dysfunction as inefficient clearance furthers inflammation and leads to adverse cardiac remodeling^46^. In accordance with our data, the “don’t eat me” signal CD47^33^, which prevents macrophage efferocytosis, is elevated in atherosclerotic plaques, and in the heart after myocardial infarction. This molecular defect is targetable as treatment with an anti-CD47 antibody improves cardiac function in multiple preclinical models of CVD, including myocardial infarction and doxorubicin cardiomyopathy^34,47^. Macrophage efferocytosis may also maintain cardiomyocyte metabolic health as dysfunctional mitochondria are exported as exophers and processed by resident macrophages^48^. Improper elimination of exophers activates inflammasomes, halts autophagy, and induces cardiac metabolic impairments and dysfunction^48^. Thus, impaired efferocytosis may also contribute to mitochondrial defects in RV cardiomyocytes, demonstrating crosstalk between these two important molecular phenotypes.

An additional pathway that could be contributing to RV failure is the ribotoxic stress response. Ribosome stress, such as damage, collisions, or stalling, triggers the RSR through ZAKα phosphorylation^49^, which can lead to pyroptosis via NACHT, LRR, and PYD domains-containing protein 1 (NLRP1) inflammasome signaling^50–52^. Besides its role in inflammation, ZAKα also modulates cellular metabolism by negatively regulating mammalian target of rapamycin (mTOR) signaling and activating AMP-activated protein kinase (AMPK)^53^. Furthermore, ROS generated via impaired regulation of the electron transport chain can initiate the RSR^44^. Consequently, there may be considerable crosstalk between the RSR, metabolic dysregulation, and RV inflammation, key cellular phenotypes consistently identified as drivers of RV failure^6,8,9^.

In addition to these three identified pathways, our kinome analysis suggests there are kinases activated in severe RVD that could be targeted to modify inflammation and metabolism (**Supplemental Table 2**). Three kinases activated in severe RVD modulate cellular metabolism and have a therapy targeting the kinase approved or in a clinical trial: AKT1^54^, GSK3β^55^, and STK11^56^. FYN, a kinase that regulates T-cell activation^57^, is the target of an investigational therapy in clinical trials for pulmonary fibrosis. Therefore, multiple kinases could be targeted to augment RV function.

## Limitations

We acknowledge our manuscript has important limitations. First, this was an analysis of young, castrated male pigs, so biological sex and age were not evaluated. However, our data are congruent with transcriptomic analyses in humans of both sexes, demonstrating progressive impairments in mitochondrial metabolism and increased inflammatory pathway engagement in RV failure^8,9^. Our snRNAseq data suggested RV fibroblast biology was altered in our model, but we did not detect significant alterations in interstitial or perivascular fibrosis in the three experimental groups (**Supplement Figure 4**). This lack of fibrosis may be related to castration, age, or a combination of the two. This hypothesis is supported by the observation that children with dilated cardiomyopathy have less fibrosis than adults^58^, and that castrated mice have reduced RV fibrosis^59^. However, the alterations in fibroblast phenotypes could indicate they are playing other roles, such as modulating RV inflammation, an emerging role of cardiac fibroblasts^60,61^. In addition, these data are strictly descriptive, and future studies are needed to determine how interventions targeting these pathways modulate RV function. Furthermore, we only evaluated two types of post-translational modification, phosphorylation and ubiquitination, and further studies are required to determine how other post-translational modifications are dysregulated along the spectrum of RVD. Unfortunately, there is no robust algorithm to predict how distinct phosphatases modulate cardiac biology, and thus harnessing our phosphatome data will require additional experiments to determine whether these changes are pathological, compensatory, or non-contributory. Finally, there were instances in which our transcriptomic and proteomic data gave divergent responses. This may be due to differences in RNA stability, translational efficiency, and proteostasis.

## Author Contributions

WT, MTL, JPC, FK, and KWP performed PAB and invasive hemodynamic procedures, developed cMRI protocol, and analyzed cMRI data. MK, RMR, RAM, and SEP performed histology experiments, collected and analyzed data. JBM, JDS, LHM isolated nuclei, developed snRNAseq pipeline and performed analysis. MK, RAM, RMR, SEP, TM, and LH performed cellular fractionations assays, performed TMT proteomics, and analyzed proteomics/phosphoproteomics data. JBM and KWP completed formal statistical analysis. JBM and KWP wrote the manuscript, which was reviewed and approved by all authors.

## Non-Standard Abbreviations

ABL1: ABL proto-oncogene 1, non-receptor tyrosine kinase
AKT1: AKT serine/threonine kinase 1
AMPK: AMP-activated protein kinase
CCL24: C-C motif chemokine ligand 24
CD47: Cluster of differentiation 47
CDK4: cyclin-dependent kinase 4
CVD: Cardiovascular disease
CXCL10: C-X-C motif chemokine ligand 10
DEG: Differentially expressed genes
EGFR: epidermal growth factor receptor
FACS: Fluorescence activated cell sorting
FAP: fibroblast activating protein
FN1: fibronectin 1
FYN: Fyn kinase
GSK3β: glycogen synthase kinase 3 beta
IFIT1: interferon-induced protein with tetratricopeptide repeats 1
ILK: integrin linked kinase
KEGG: Kyoto Encyclopedia of Genes and Genomes
LATS2: large tumor suppressor kinase 2
LONP1: lon peptidase 1
LRRK2: leucine rich repeat kinase 2
LYVE1: lymphatic vessel endothelial hyaluronan receptor 1
MARK1: microtubule affinity regulating kinase 1
mitoUPR: mitochondrial unfolded protein response
MRC1: mannose receptor C type 1
mTORC1: mammalian target of rapamycin complex 1
NLRP1: NACT, LRR, and PYD domains-containing protein 1
NTRK1: neurotrophic receptor tyrosine kinase 1
OCT: optimal cutting temperature
PDGFRβ: platelet derived growth factor receptor beta
PDK3: pyruvate dehydrogenase kinase 3
PHKG2: phosphorylase kinase catalytic subunit gamma 2
PLAC8: placenta associated 8
POSTN: periostin
RAF1: raf-1 proto-oncogene serine/threonine-protein kinase
RIOK1: RIO kinase 1
RVD: Right ventricular dysfunction
STK11: serine/threonine kinase 11
TBK1: TANK-binding kinase 1
TCA: Tricarboxylic acid
TTN: titin
ZAKα: sterile alpha motif and leucine zipper containing kinase AZK

---

**Supplemental Figure 1:**
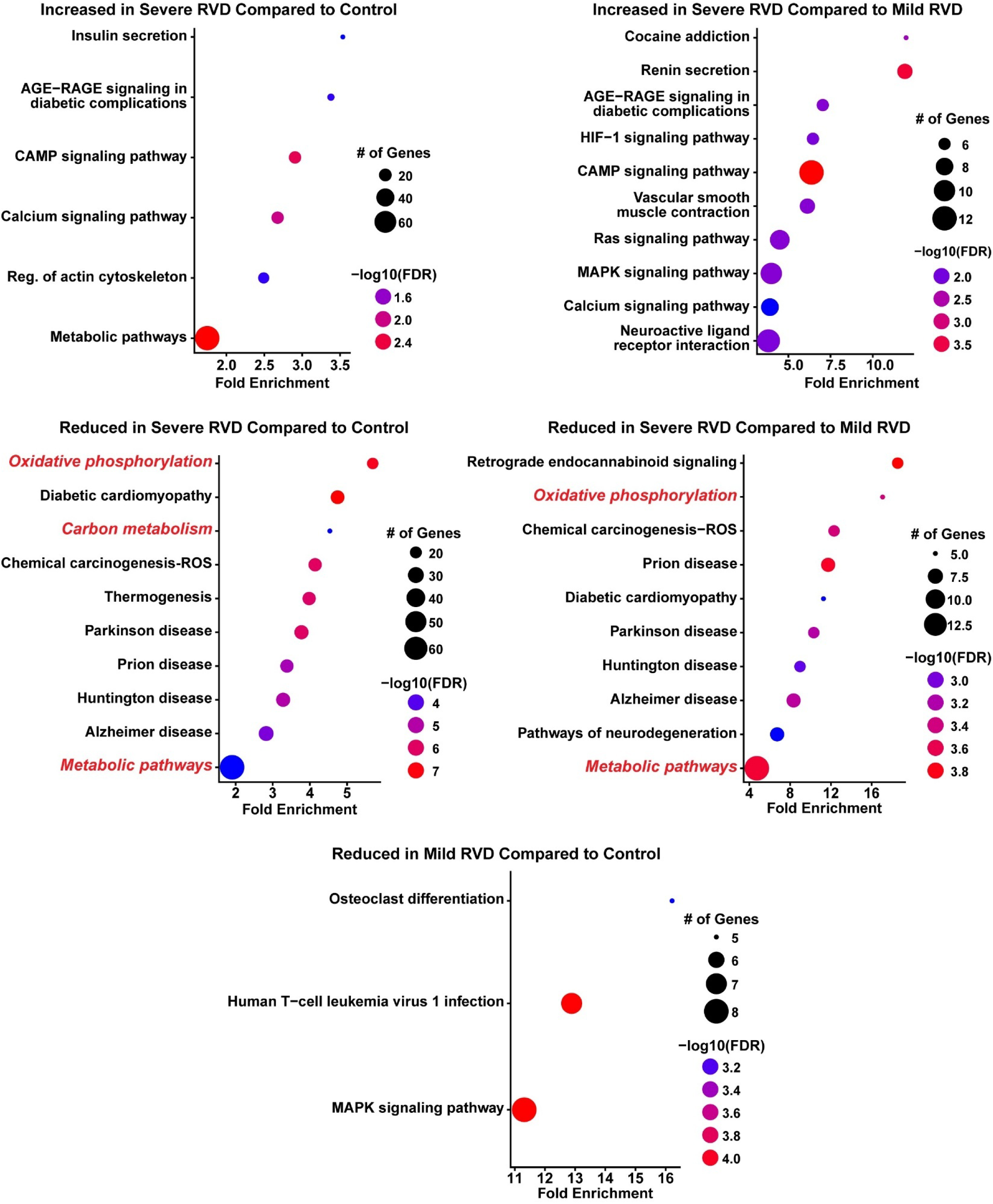
snRNAseq identifies metabolic derangements in progressive RV failure. Pathway analysis of differentiallly expressed cardiomyocyte genes identified metabolic pathways are reduced only in severe RVD (red italics).

**Supplemental Figure 2:**
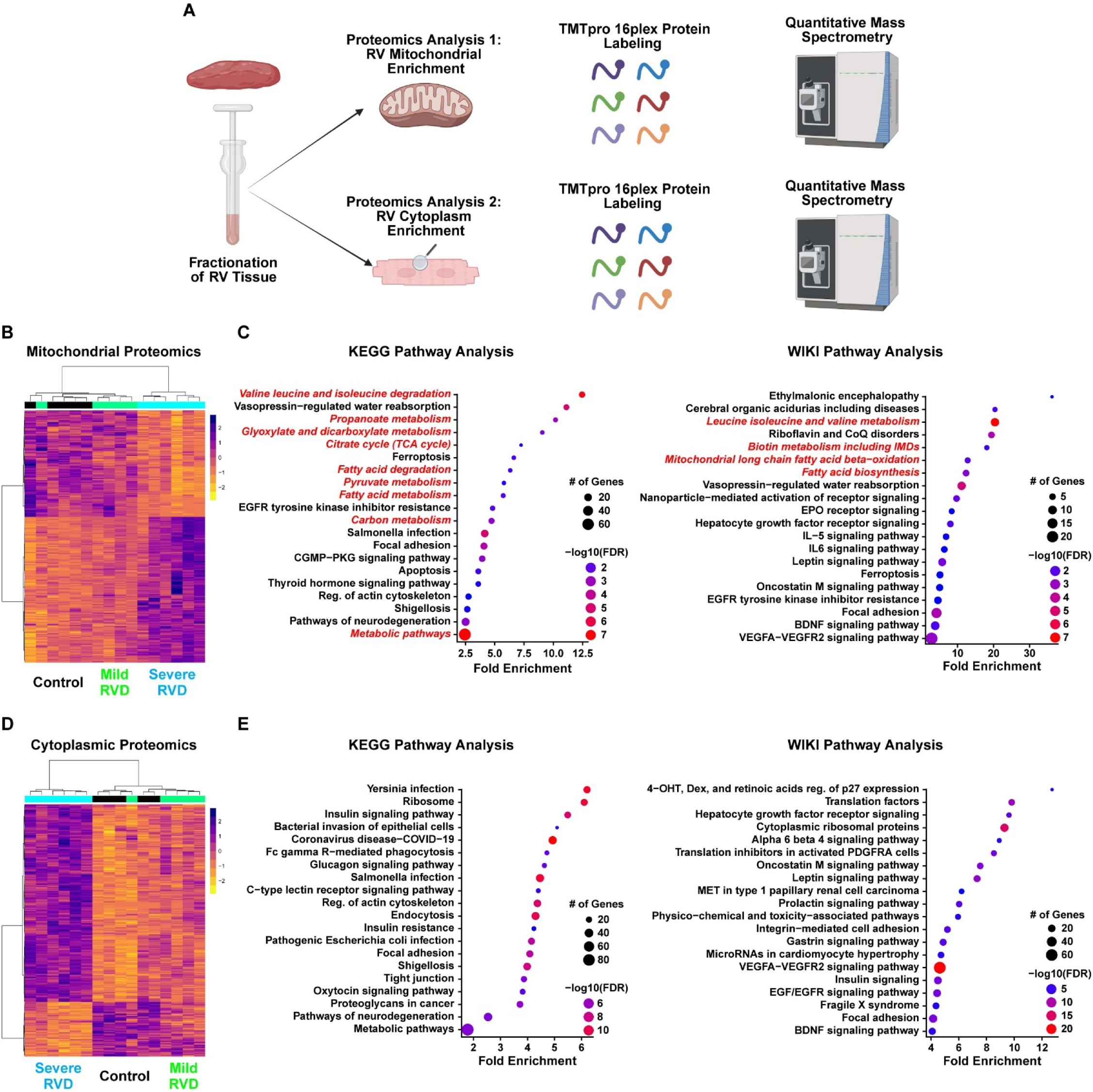
Proteomics approaches and pathways enriched with progressive RV failure. (A) Schematic of mitochondrial and cytoplasmic proteomics approaches. (B) Hierarchical cluster analysis of mitochondrial proteomics cluster control with mild RVD. Pathway analysis using (C) Kyoto Encyclopedia of Genes and Genomes (KEGG) and (D) Wiki pathway databases identified disrupted metabolic pathways (red italics). (D) Hierarchical cluster analysis of the cytoplasmic proteomics group cluster mild RVD and control together. Pathway analysis with (E) KEGG and (F) WIKI databases did not identify altered metabolic pathways in the cytoplasm.

**Supplemental Figure 3:**
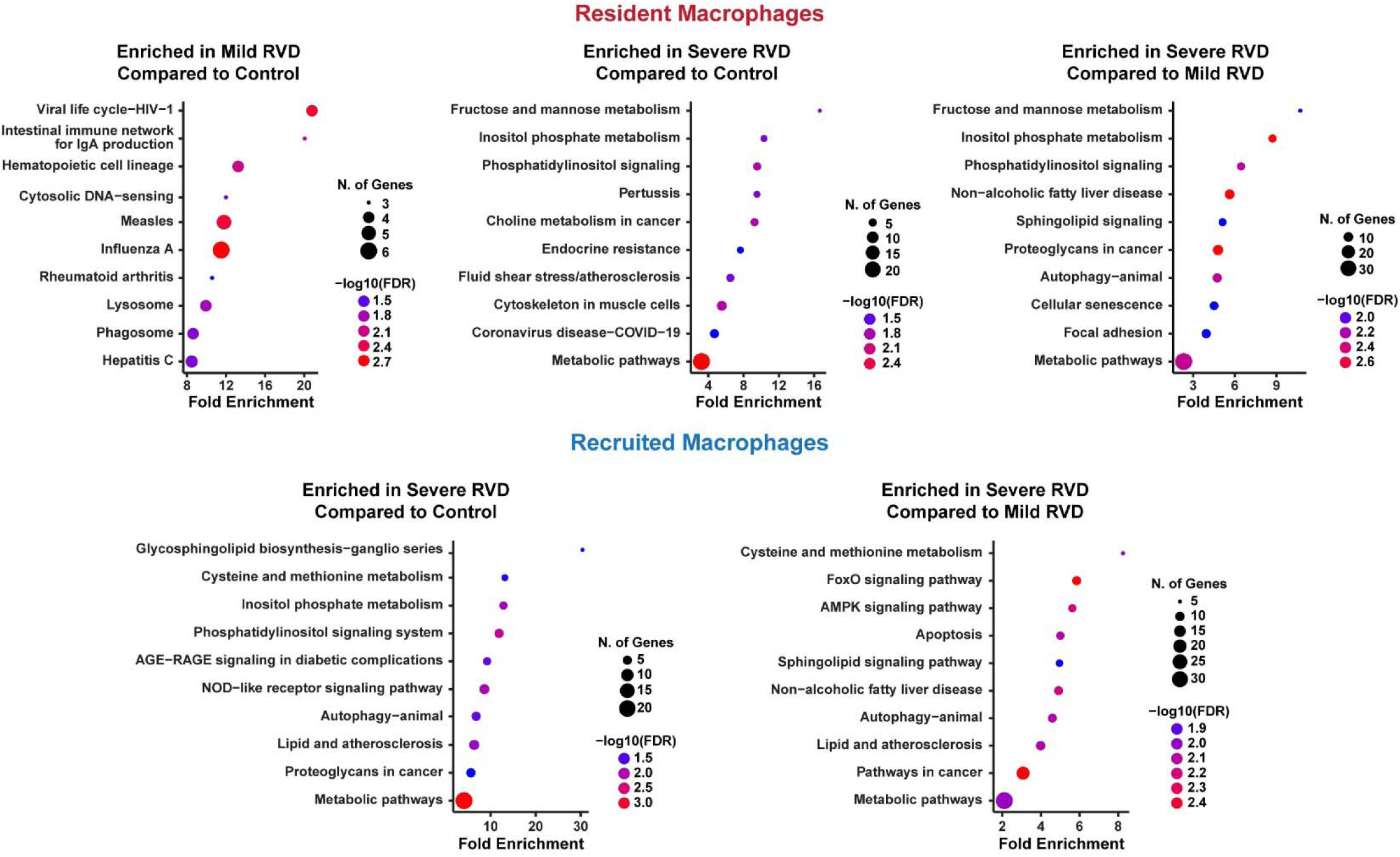
Pathways upregulated in resident and recruited macrophages in pigs with mild or severe RVD. Inflammatory and metabolic pathways were enriched in both resident and recruited macrophages.

**Supplemental Figure 4:**
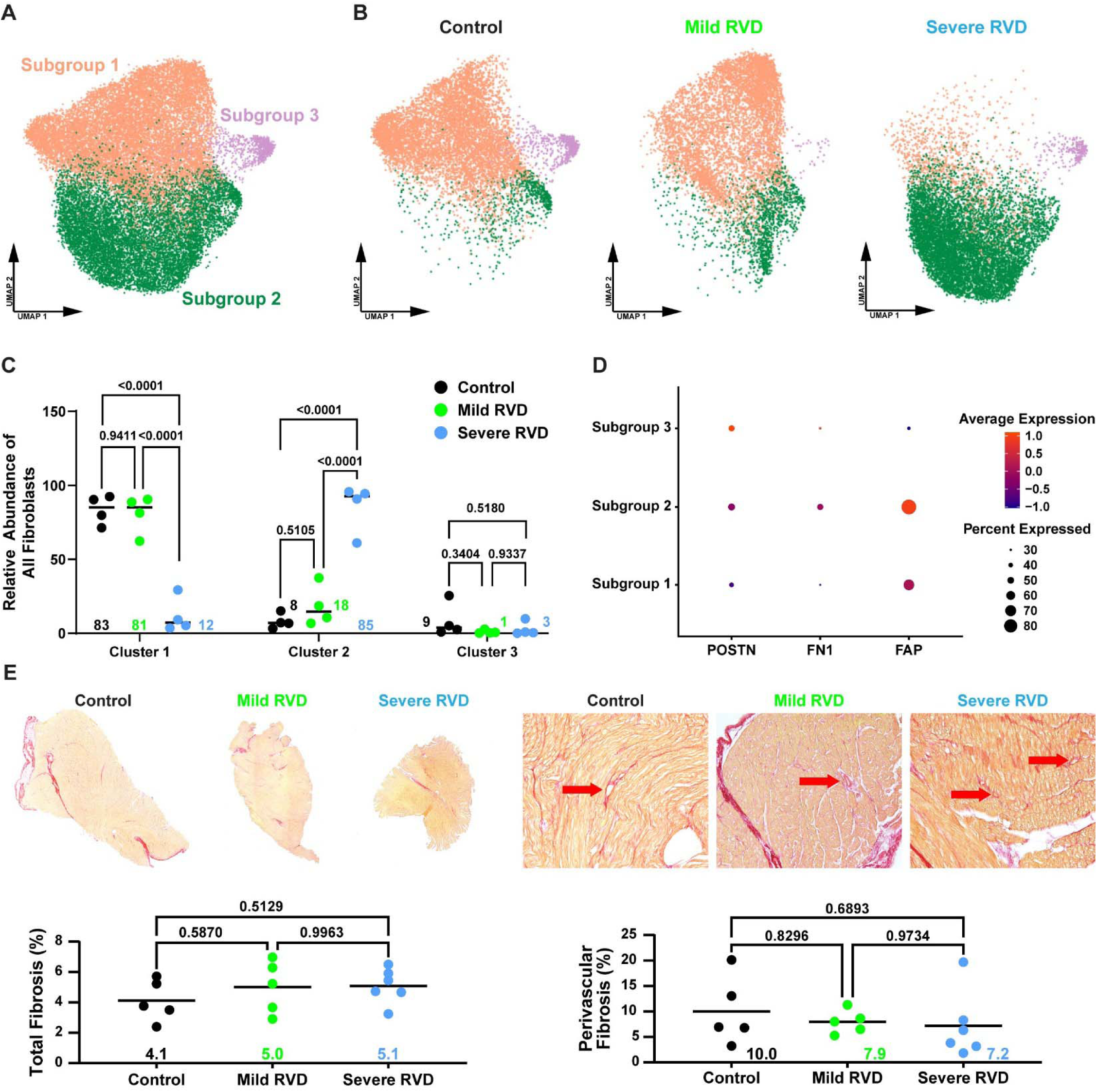
snRNAseq identifies alterations in fibroblast regulation in progressive RV Failure. (A) UMAP visualization of fibroblast subtypes (B) split by experimental group. (C) Cluster 1 primarily contained nuclei from control and mild RVD, while cluster 2 was mainly severe RVD. (D) Cluster 2 contained nuclei with elevated expression of genes associated with fibroblast activation. (E) Analysis of total fibrosis and perivascular fibrosis revealed no changes in fibrosis between groups (red arrows: perivascular fibrosis). POSTN: periostin, FN1: fibronectin 1, FAP: fibroblast activated protein.

**Supplemental Figure 5:**
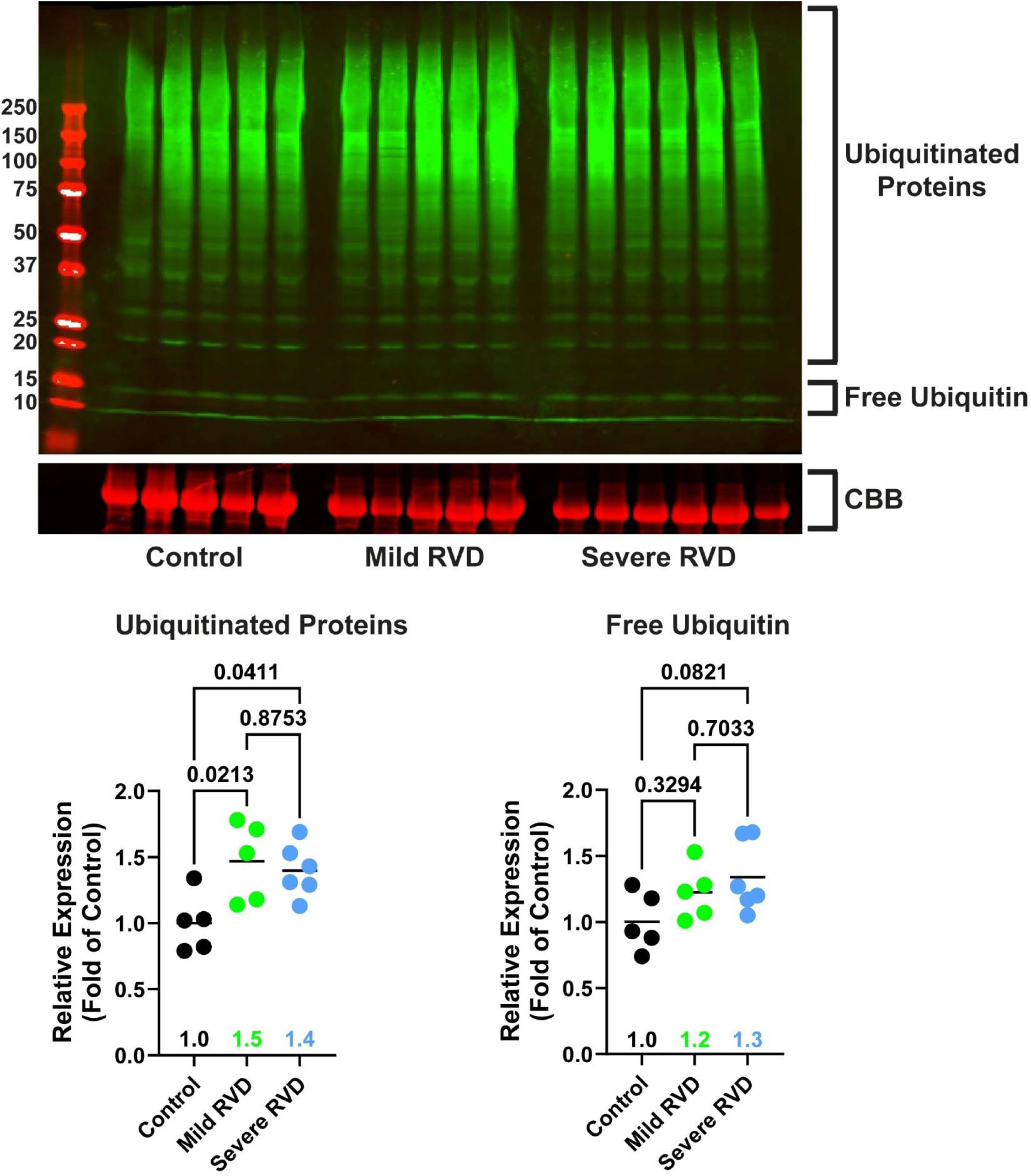
Protein ubiquitination. Protein ubiquitination was similarly elevated in mild RVD and severe RVD compared to control.

**Supplemental Figure 6:**
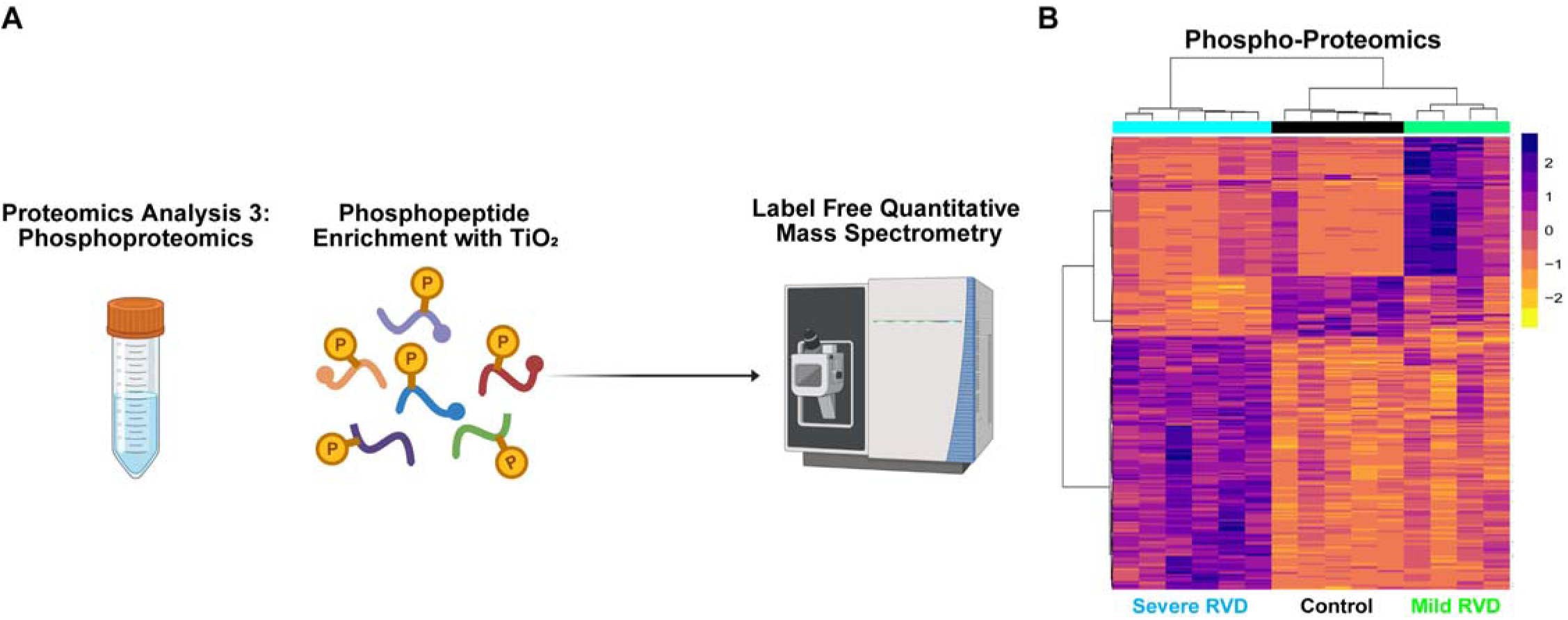
Phosphoproteomics approach. (A) Schematic of phosphoproteomics analysis. (B) Mild RVD samples clustered with the control group on hierarchical cluster analysis.

**Supplemental Table 1:**
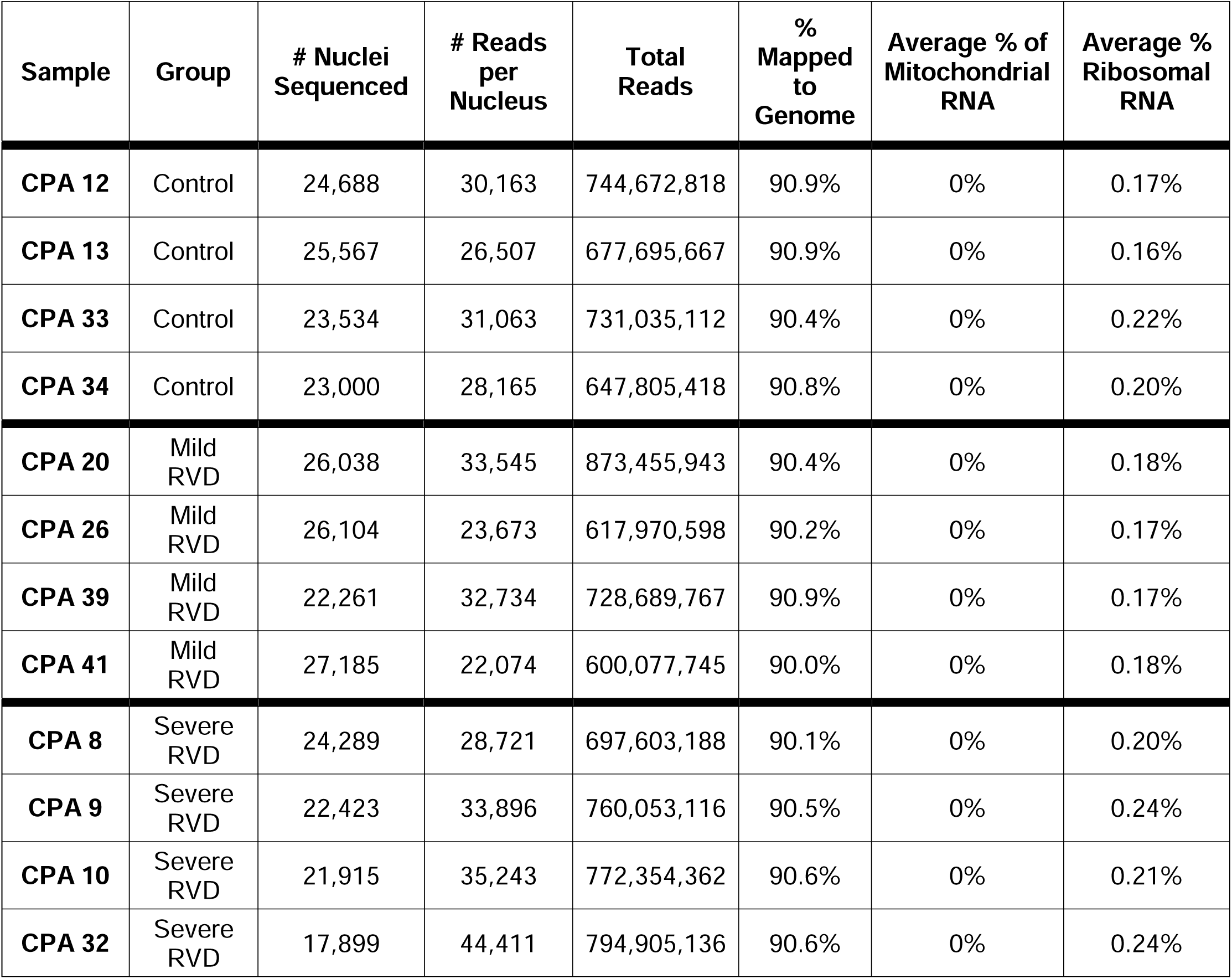
snRNAseq summary.

**Supplemental Table 2:**
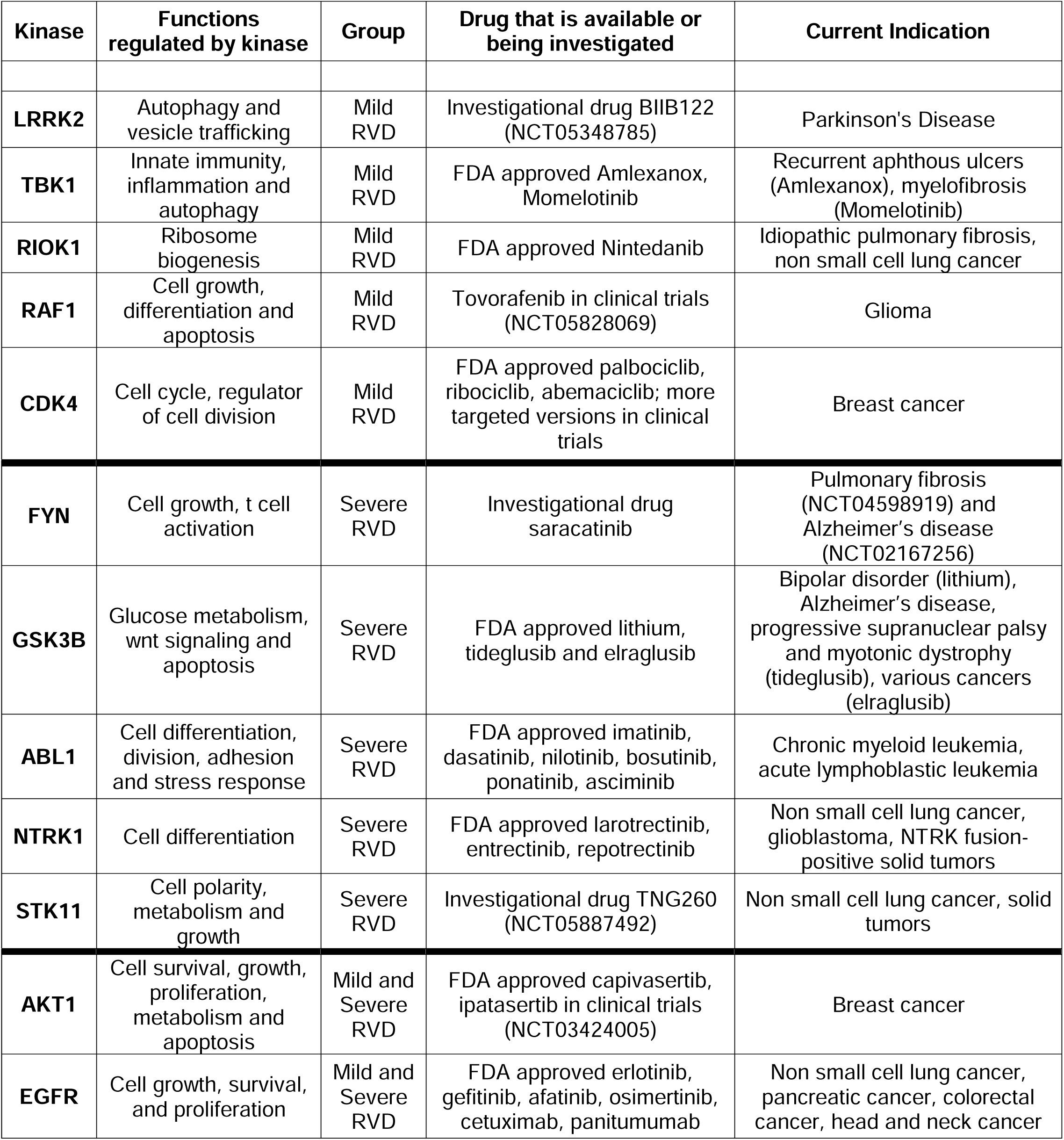
Kinases elevated in mild and severe RVD with approved or investigational therapies.

