## Supplemental Data for "Multimodality Molecular Profiling Nominates Targetable Mechanisms in Progressive RV Dysfunction"

**Supplemental Table 1: snRNAseq summary.**

| **Sample** | **Group** | **# Nuclei Sequenced** | **# Reads per Nucleus** | **Total Reads** | **% Mapped to Genome** | **Average % of Mitochondrial RNA** | **Average % Ribosomal RNA** |
| --- | --- | --- | --- | --- | --- | --- | --- |
| **CPA 12** | Control | 24,688 | 30,163 | 744,672,818 | 90.9% | 0% | 0.17% |
| **CPA 13** | Control | 25,567 | 26,507 | 677,695,667 | 90.9% | 0% | 0.16% |
| **CPA 33** | Control | 23,534 | 31,063 | 731,035,112 | 90.4% | 0% | 0.22% |
| **CPA 34** | Control | 23,000 | 28,165 | 647,805,418 | 90.8% | 0% | 0.20% |
| **CPA 20** | Mild RVD | 26,038 | 33,545 | 873,455,943 | 90.4% | 0% | 0.18% |
| **CPA 26** | Mild RVD | 26,104 | 23,673 | 617,970,598 | 90.2% | 0% | 0.17% |
| **CPA 39** | Mild RVD | 22,261 | 32,734 | 728,689,767 | 90.9% | 0% | 0.17% |
| **CPA 41** | Mild RVD | 27,185 | 22,074 | 600,077,745 | 90.0% | 0% | 0.18% |
| **CPA 8** | Severe RVD | 24,289 | 28,721 | 697,603,188 | 90.1% | 0% | 0.20% |
| **CPA 9** | Severe RVD | 22,423 | 33,896 | 760,053,116 | 90.5% | 0% | 0.24% |
| **CPA 10** | Severe RVD | 21,915 | 35,243 | 772,354,362 | 90.6% | 0% | 0.21% |
| **CPA 32** | Severe RVD | 17,899 | 44,411 | 794,905,136 | 90.6% | 0% | 0.24% |

**Supplemental Figure 1: snRNAseq identifies metabolic derangements in progressive RV failure.** Pathway analysis of differentiallly expressed cardiomyocyte genes identified metabolic pathways are reduced only in severe RVD (red italics).

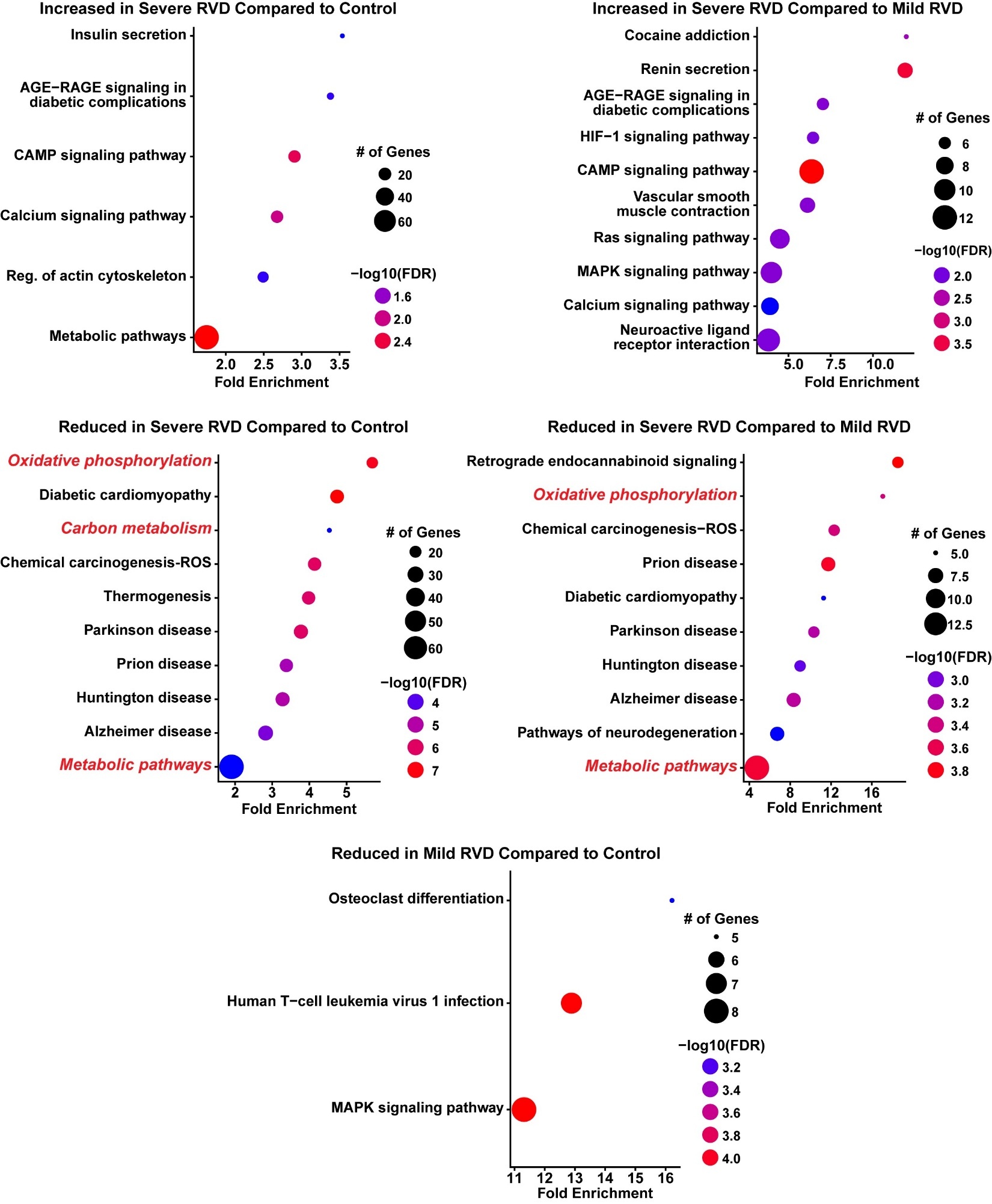

**Supplemental Figure 2**: **Proteomics approaches and pathways enriched with progressive RV failure**. (A) Schematic of mitochondrial and cytoplasmic proteomics approaches. (B) Hierarchical cluster analysis of mitochondrial proteomics cluster control with mild RVD. Pathway analysis using (C) Kyoto Encyclopedia of Genes and Genomes (KEGG) and (D) Wiki pathway databases identified disrupted metabolic pathways (red italics). (D) Hierarchical cluster analysis of the cytoplasmic proteomics group cluster mild RVD and control together. Pathway analysis with (E) KEGG and (F) WIKI databases did not identify altered metabolic pathways in the cytoplasm.

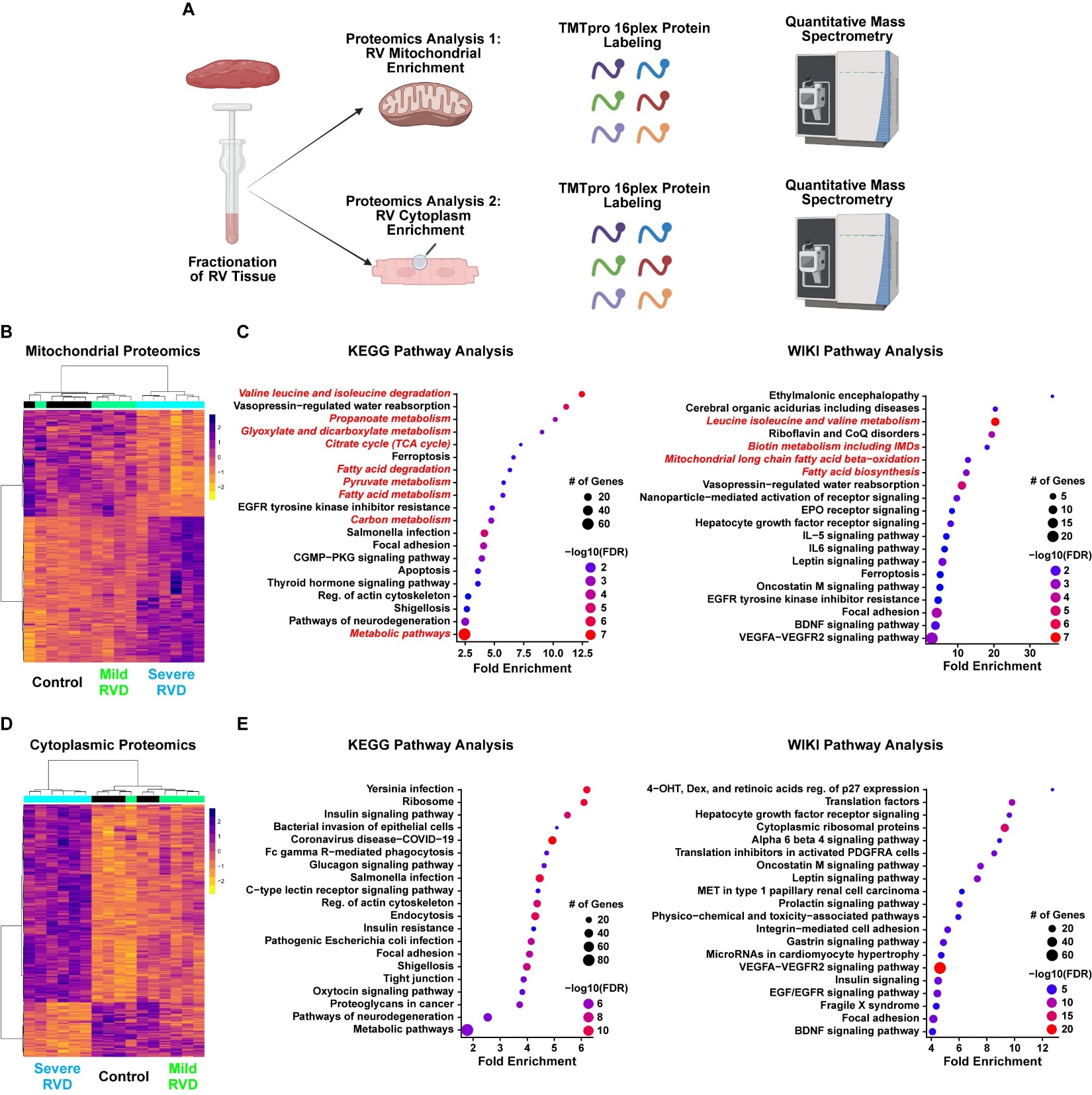

**Supplemental Figure 3: Pathways upregulated in resident and recruited macrophages in pigs with mild or severe RVD**. Inflammatory and metabolic pathways were enriched in both resident and recruited macrophages.

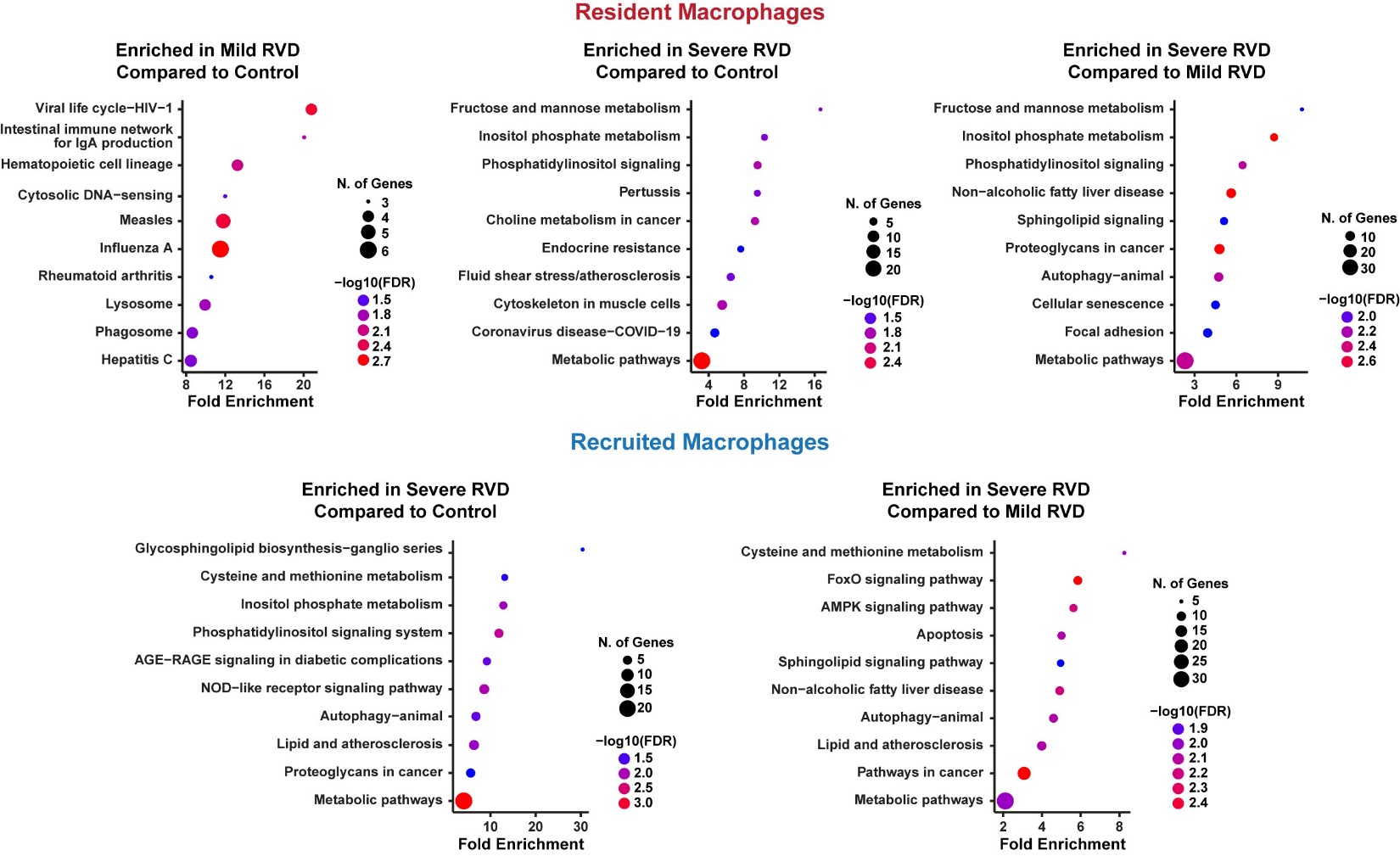

**Supplemental Figure 4: snRNAseq identifies alterations in fibroblast regulation in progressive RV Failure**. (A) UMAP visualization of fibroblast subtypes (B) split by experimental group. (C) Cluster 1 primarily contained nuclei from control and mild RVD, while cluster 2 was mainly severe RVD. (D) Cluster 2 contained nuclei with elevated expression of genes associated with fibroblast activation. (E) Analysis of total fibrosis and perivascular fibrosis revealed no changes in fibrosis between groups (red arrows: perivascular fibrosis). POSTN: periostin, FN1: fibronectin 1, FAP: fibroblast activated protein.

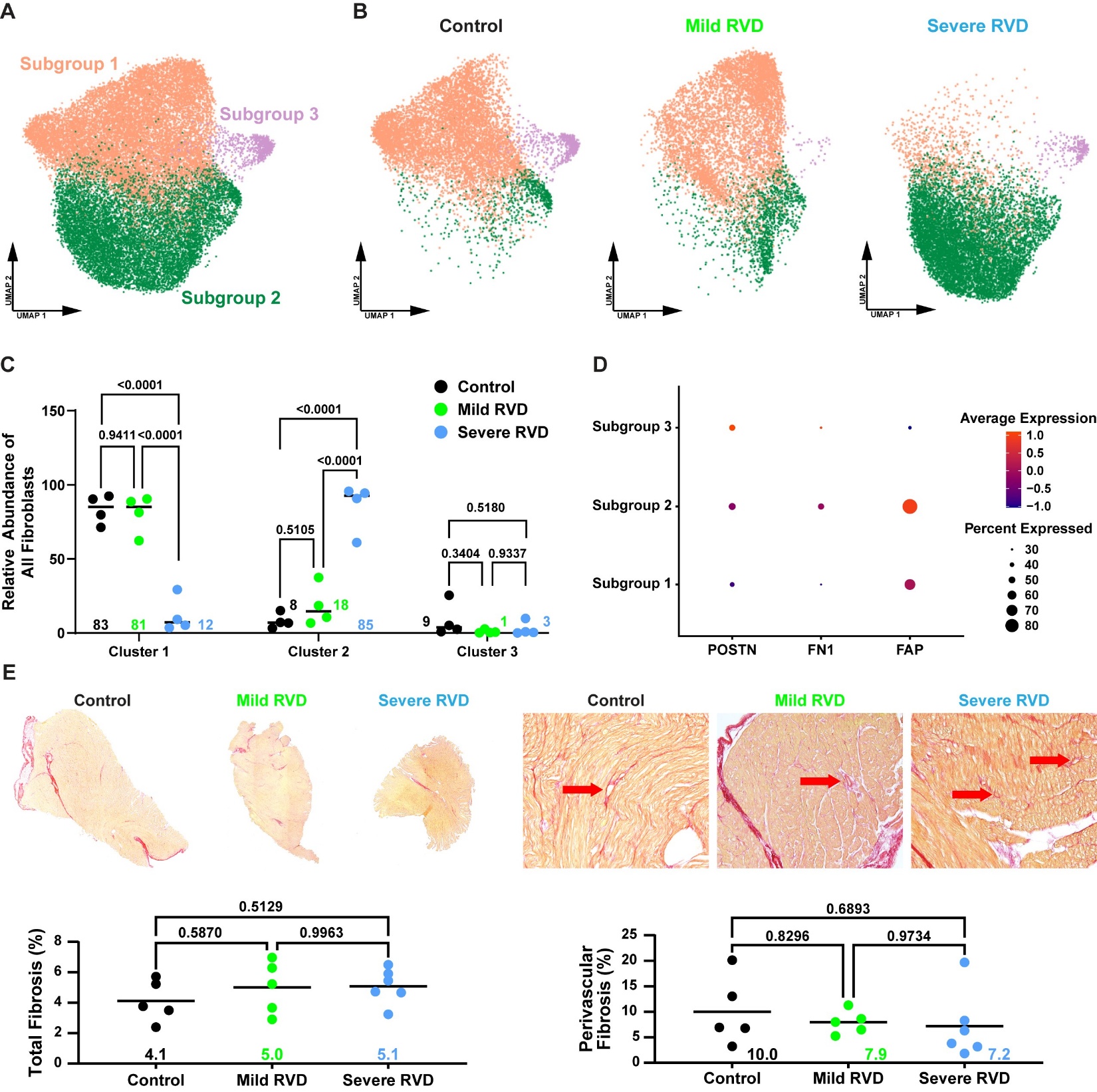

**Supplemental Figure 5: Protein ubiquitination.** Protein ubiquitination was similarly elevated in mild RVD and severe RVD compared to control.

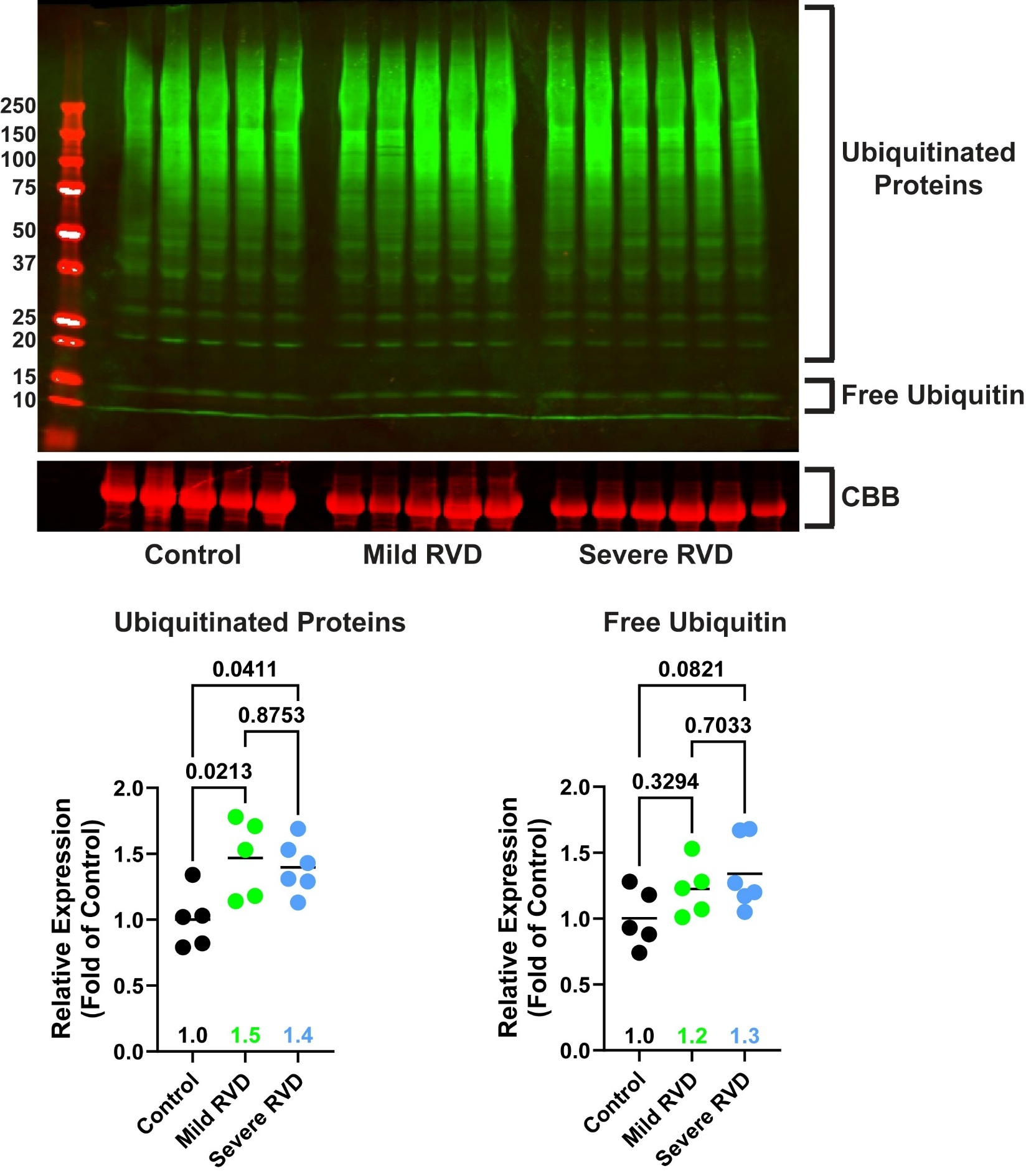

**Supplemental Figure 6: Phosphoproteomics approach.** (A) Schematic of phosphoproteomics analysis. (B) Mild RVD samples clustered with the control group on hierarchical cluster analysis.

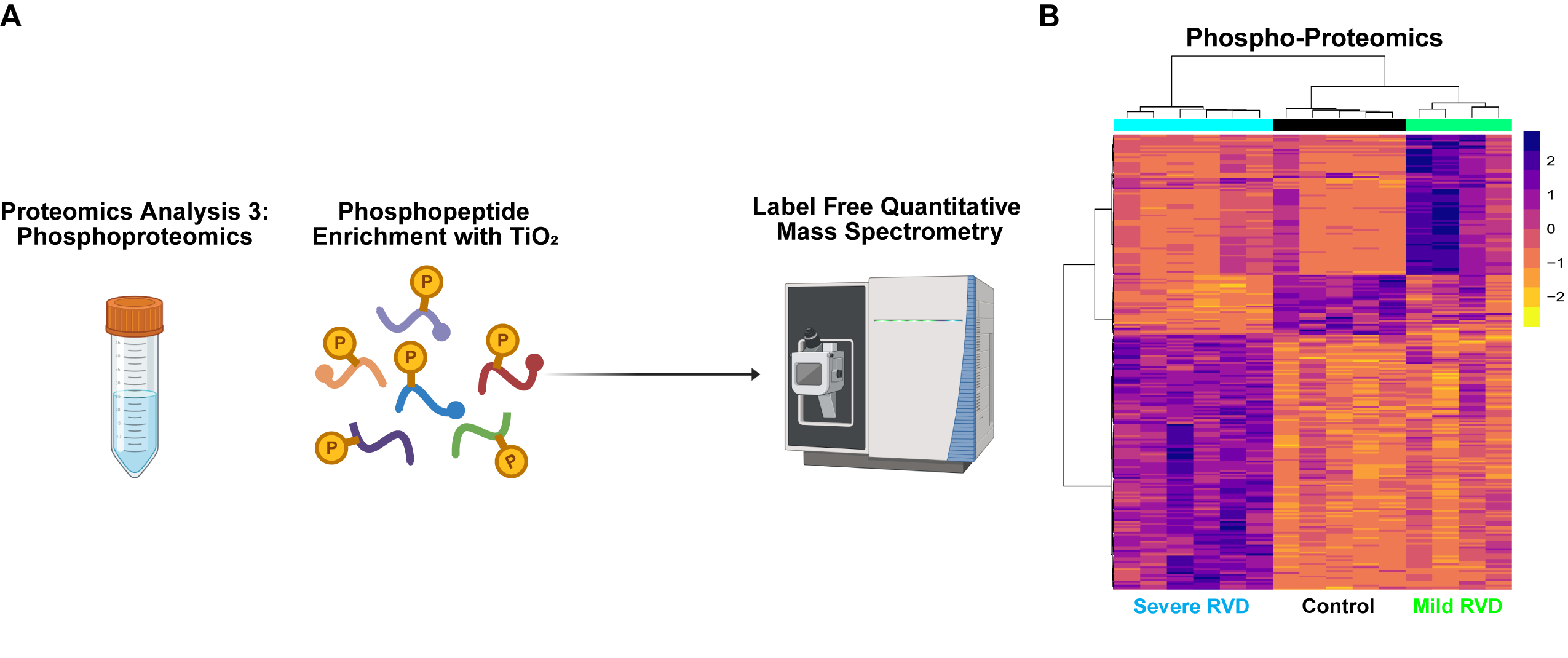

**Supplemental Table 2: Kinases elevated in mild and severe RVD with approved or investigational therapies.**

| **Kinase** | **Functions regulated by kinase** | **Group** | **Drug that is available or being investigated** | **Current Indication** |
| --- | --- | --- | --- | --- |
| **LRRK2** | Autophagy and vesicle trafficking | Mild RVD | Investigational drug BIIB122 (NCT05348785) | Parkinson's Disease |
| **TBK1** | Innate immunity, inflammation and autophagy | Mild RVD | FDA approved Amlexanox, Momelotinib | Recurrent aphthous ulcers (Amlexanox), myelofibrosis (Momelotinib) |
| **RIOK1** | Ribosome biogenesis | Mild RVD | FDA approved Nintedanib | Idiopathic pulmonary fibrosis, non small cell lung cancer |
| **RAF1** | Cell growth, differentiation and apoptosis | Mild RVD | Tovorafenib in clinical trials (NCT05828069) | Glioma |
| **CDK4** | Cell cycle, regulator of cell division | Mild RVD | FDA approved palbociclib, ribociclib, abemaciclib; more targeted versions in clinical trials | Breast cancer |
| **FYN** | Cell growth, t cell activation | Severe RVD | Investigational drug saracatinib | Pulmonary fibrosis (NCT04598919) and Alzheimer’s disease (NCT02167256) |
| **GSK3B** | Glucose metabolism, wnt signaling and apoptosis | Severe RVD | FDA approved lithium, tideglusib and elraglusib | Bipolar disorder (lithium), Alzheimer’s disease, progressive supranuclear palsy and myotonic dystrophy (tideglusib), various cancers (elraglusib) |
| **ABL1** | Cell differentiation, division, adhesion and stress response | Severe RVD | FDA approved imatinib, dasatinib, nilotinib, bosutinib, ponatinib, asciminib | Chronic myeloid leukemia, acute lymphoblastic leukemia |
| **NTRK1** | Cell differentiation | Severe RVD | FDA approved larotrectinib, entrectinib, repotrectinib | Non small cell lung cancer, glioblastoma, NTRK fusion-positive solid tumors |
| **STK11** | Cell polarity, metabolism and growth | Severe RVD | Investigational drug TNG260 (NCT05887492) | Non small cell lung cancer, solid tumors |
| **AKT1** | Cell survival, growth, proliferation, metabolism and apoptosis | Mild and Severe RVD | FDA approved capivasertib, ipatasertib in clinical trials (NCT03424005) | Breast cancer |
| **EGFR** | Cell growth, survival, and proliferation | Mild and Severe RVD | FDA approved erlotinib, gefitinib, afatinib, osimertinib, cetuximab, panitumumab | Non small cell lung cancer, pancreatic cancer, colorectal cancer, head and neck cancer |
